# A fetal wave of human type-3 γδ T cells with restricted TCR diversity persists into adulthood

**DOI:** 10.1101/2020.08.14.248146

**Authors:** Likai Tan, Alina Suzann Fichtner, Anja Bubke, Ivan Odak, Christian Schultze-Florey, Christian Koenecke, Reinhold Förster, Michael Jarek, Constantin von Kaisenberg, Alina Borchers, Ulf Panzer, Christian Krebs, Sarina Ravens, Immo Prinz

**Affiliations:** Institute of Immunology, Hannover Medical School, Hannover, Germany; Department of Hematology, Hemostasis, Oncology and Stem Cell Transplantation, Hannover Medical School, Hannover, Germany; Genome Analytics, Helmholtz Centre for Infection Research, Braunschweig, Germany; Department of Obstetrics, Gynecology and Reproductive Medicine, Hannover Medical School, Hannover, Germany; Translational Immunology, III. Department of Medicine, Hamburg Center for Translational Immunology (HCTI), University Medical Center Hamburg-Eppendorf, Hamburg, Germany; Cluster of Excellence RESIST (EXC 2155), Hannover Medical School, Hannover, Germany

**Keywords:** Human γδ T cells, paired γδ TCR sequencing, single-cell RNA sequencing, γδ T cell development, γδT17 cells

## Abstract

Accumulating evidence suggests that the human embryonic thymus produces distinct waves of innate effector γδ T cells. However, it is unclear whether this process comprises a dedicated subset of IL-17-producing γδ T (γδT17) cells, like reported in mice. Here we present a novel protocol for high-throughput paired γδ TCR-sequencing, which in combination with single-cell RNA-sequencing revealed a high heterogeneity of effector γδ T cell clusters. While immature γδ T cell clusters displayed mixed and diverse TCR, effector cell types in neonatal and adult blood segregated according to γδTCR usage. In adult samples, mature Vδ1^+^ T cells segregated into exhausted PD-1^hi^ and active PD-1^low^ clusters. Among Vγ9Vδ2^+^ T cell subsets, we identified distinct PLZF-positive effector γδ T cell clusters with innate type-1 and type-3 T cell signatures that were already detectable in a public dataset of early embryonic thymus organogenesis. Together, this suggests that functionally distinct waves of human innate effector γδ T cells including CCR6^+^ γδT17 cells develop in the early fetal thymus and persist into adulthood.

## Introduction

γδ T cells are an evolutionary conserved subset of T lymphocytes that can respond to microbial stimuli and provide tissue-surveillance independent of MHC-peptide recognition. Due to their seemingly intrinsic cytotoxicity towards a large array of tumors, γδ T cells are the subject of many current efforts to design anti-cancer immunotherapies^1^. Although generally regarded as rather innate T cells, γδ T cells display a great level of phenotypic heterogeneity^2^.

The most abundant γδ T lymphocytes in human adult blood are defined by a semi-invariant T cell receptor (TCR) composed of a TCRγ chain (*TRG*) using the variable (V) segment Vγ9 (*TRGV9*) rearranged to the joining (J) segment JP (*TRGJP*), and a TCRδ chain (*TRD*) using a Vδ2 segment^3^. Such Vγ9Vδ2^+^ T cells still display a considerable TCR diversity, but collectively respond to the intracellular accumulation of microbial and tumor-derived metabolites called phosphoantigens^4^, which are somehow support binding of canonical Vγ9Vδ2^+^ T cells to butyrophilin-molecules BTN3A1^5^ and BTN2A1^6,7^. Furthermore, Vγ9Vδ2^+^ T cells share common features with other human innate-like T cells like invariant natural killer T (NKT) and mucosa-associated invariant T (MAIT) cells, including expression of PLZF^8^. The remaining non-Vγ9Vδ2^+^ human γδ T cell subsets express diverse *TRG* rearrangements, preferentially paired with Vδ1^3^. Information about specific ligands binding to Vδ1 ^+^ TCRs is still fragmentary, however, recent findings showed expansion of individual Vδ1^+^ clones in response to environmental, e.g. viral stimuli^9,10^.

In this setting, the correlation of γδTCR usage, TCR-specific activation and acquisition of γδ T cell effector function is in the central focus of the field. In mice, distinct waves of effector γδ T cells with restricted TCR diversity develop sequentially in the fetal thymus^11^. In particular, IL-17-producing effector γδ T (γδT17) cells are exclusively generated before birth and persist throughout the entire life as self-renewing, tissue-resident T cells in a diverse range of tissues^12,13^. There are parallels in humans, as Vγ9Vδ2^+^ T cells with restricted TCR diversity are preferentially generated before birth, expand postnatally and persist into adulthood^14–17^. However, if and to which extent human γδ T cells acquire effector function already in the thymus is a matter of debate. Postnatal thymi contained γδ T cells with immense TCR diversity^18^, but these were not equipped with effector functions such as IFN-γ or IL-17 production^19^. In contrast, the human fetal thymus generates invariant effector γδ T cells that express invariant germline-encoded CDR3γ and CDR3δ repertoires^20^. Furthermore, a recent study compared human Vδ1^+^ and Vδ2^+^ γδ T cells by single-cell RNA sequencing (scRNA-seq) and concluded that both displayed parallel maturation trajectories, starting with expression of maturation-defining genes including *CCR7, IL7R*, and *CD27* and gradually acquiring cytotoxicity-related genes such as *NKG7, PRF1*, and *FCGR3A*^21^.

To directly assess the role of the TCR in guiding the activation and differentiation of individual γδ T cell clones, we sorted total γδ T cells from cord and adult blood and combined scRNA-seq with single-cell sequencing of paired *TRG* and *TRD* rearrangements (scTCR-seq). We found a functional heterogeneity of immature and effector γδ T cells, that segregated according to γδTCR usage. Notably, we identified distinct γδT17 and Th1-like and cytotoxic effector (γδT1) subsets of Vγ9Vδ2^+^ T cells that were already present in neonatal blood and provide evidence that these might be exclusively generated in early fetal waves. In addition, we established a minimal flow cytometric panel comprising three TCR-specific and nine surface markers that reproduces the phenotypic map of human γδ T cell subsets obtained by scRNA-seq.

## Results

### scRNA-seq reveals high heterogeneity among human γδ T cells

To investigate transcriptional programs of human γδ T cells at birth and in adulthood, total γδ T cells were isolated from two unrelated neonatal cord blood samples (CB_donor1 and CB_donor2) and two unrelated adult peripheral blood samples (PB_donor1 and PB_donor2) via flow cytometric sorting for scRNA-seq (**Supplementary Fig. 1**). After quality control, a total of 23,760 γδ T cells from all respective donors (median gene number = 1,307; median counts = 3,266) were considered for post-analysis (**Supplementary Fig. 2a**). Based on transcriptomic profiles, these γδ T cells were projected by Uniform Manifold Approximation and Projection (UMAP) and unsupervised clustering identified 12 cell clusters (c1 – c12) (**Fig. 1a**). A polarized age-dependent distribution was observed: Neonatal γδ T cells dominated c1 – c5 and adult γδ T cells formed the large majority of c10 – c12 (**Fig. 1b**). Clusters in the middle of the UMAP (c6 – c9) were more heterogeneous. Notably, 90.2% of c5 cells were from CB_donor1, and 96.2% of the large cluster c11 originated from PB_donor2, indicating variances of phenotypes on the individual’s level (**Fig. 1c**). However, all four donors actually contributed to all 12 clusters (**Supplementary Fig. 2b**). The map of the Top-10 differentially expressed genes (DEGs) in each cluster demonstrates the distinct gene expression profiles of each cluster (**Fig. 1d**). Among the 1,934 DEGs, we found T cell-related regulatory transcription factors (TFs), cytokine receptors, cytotoxic genes, T cell activation genes, and natural killer (NK) cell receptors (**Supplementary Fig. 2c**).

**Figure 1:**
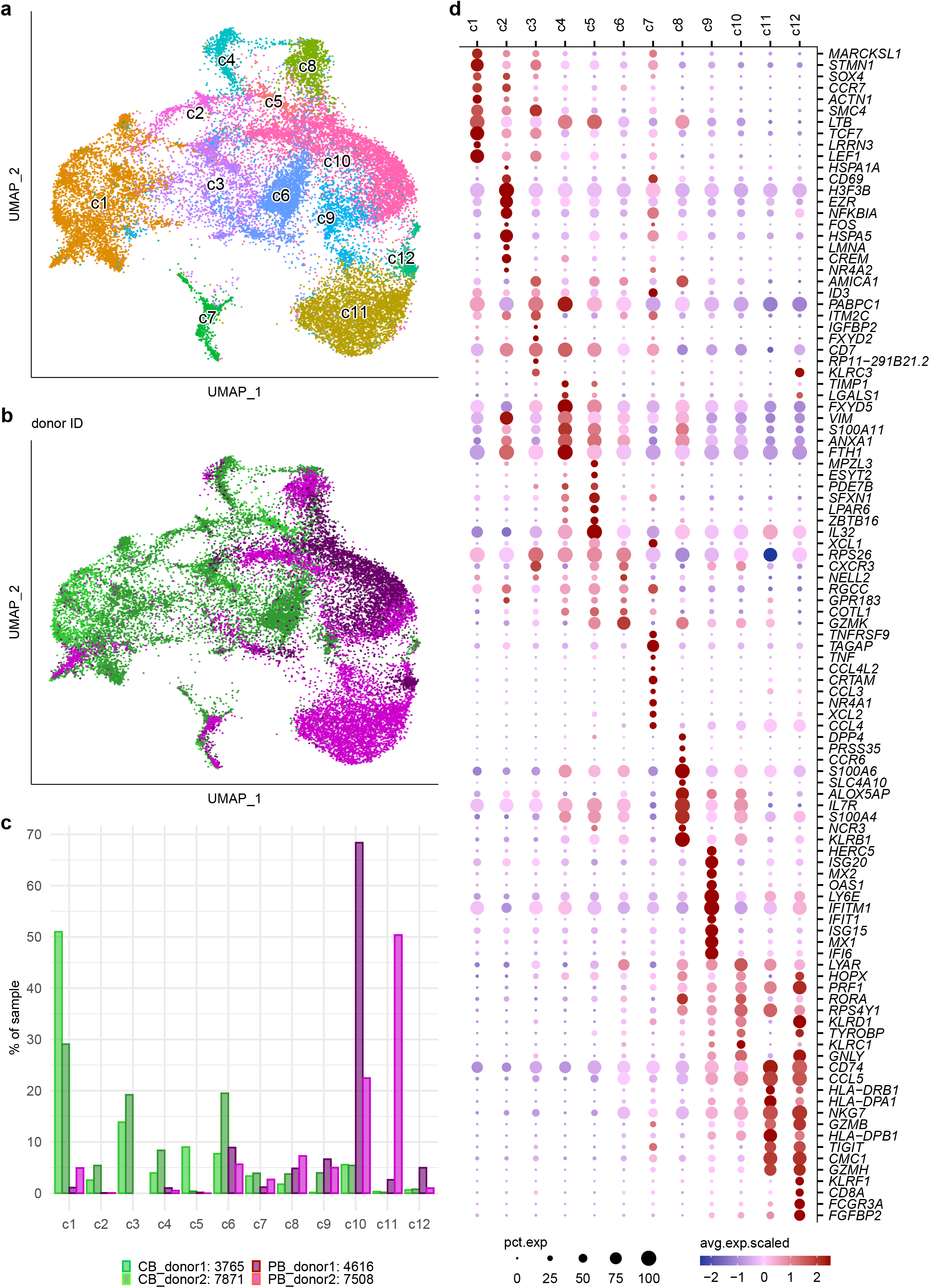
scRNA-seq reveals high heterogeneity among human γδ T cells. Single-cell transcriptome libraries were generated from FACS-sorted γδ T cells of neonatal cord blood (CB_donor1 and CB_donor2; n = 2) or adult peripheral blood (PB_donor1 and PB_donor2; n = 2). (**a,b**) Uniform Manifold Approximation and Projection (UMAP) was adopted for visualization of cells. Each point represents a single cell. (**a**) Individual cells of all four donors are colored by clusters (c1-c12) that were obtained from un-supervised clustering. (**b**) Individual cells are color-coded by the respective four donors. (**c**) Bar plot reveals fractions of absolute cell numbers that contributed from each donor to c1-c12. Total cell numbers of donors that were considered for analysis are indicated in the figure legend. (**d**) Dot plot shows the Top-10 upregulated differentially expressed genes (DEGs, rows) per cluster. Each column reflects a single cell, the color bars above represent c1-c12. For visualization, cell numbers were downsampled to max. 800 cells per cluster. Gene expression values were scaled to a Log2-fold change (logFC). DEGs were defined by being expressed on more than 10% of cells in at least one cluster (min.pct ≥ 10%), with absolute logFC ≥ 0.25 and adjusted p-value (adj.p) ≤ 0.01 (bimod test provided by Seurat package).

Next, we wondered how the heterogeneous gene expression patterns of the identified clusters reflected their potential functional commitment. Interestingly, a proportion of DEGs was expressed on more than one identified cluster (**Fig. 1d and Supplementary Fig. 2c**), suggesting overlapping traits. To understand how the 12 identified clusters were interwoven, all DEGs were subjected to unsupervised clustering based on their average expression per cluster. This analysis identified seven gene co-expression modules (GM) (**Fig. 2a and Supplementary Table S1**), which were annotated on the basis of gene ontology enrichment analysis (**Supplementary Fig. 3**). *Vice versa*, modular scoring revealed the differential expression of these GMs among the 12 γδ T cell clusters defined above (**Fig. 2b**). The gene module A (GM_A) represented an immature naïve state of T cell differentiation and was enriched within the majority of neonatal γδ T cells (c1, c3). GM_A included genes such as *LEF1* and *TCF7* that are key regulators of naïve T cells^22,23^ as well as known regulatory TFs of murine immature γδ T cells (e.g. *SOX4*)^13,24^ (**Fig. 2a**). The innate T cell differentiation gene module GM_B was present in a variety of γδ T cell clusters (c4, c5, c6, c8, and c10) and comprised the TFs GATA3 and *ZBTB16* (PLZF), described to be critical for the development of invariant NKT cells and mucosal-associated T cells^25,26^. GM_C represented cell proliferation-associated genes in c2, while GM_E was highly enriched in interferon (IFN) type I signaling-related genes in c9. The module GM_G indicated acute T cell activation in c2 and c7. Of note, the type-3 immunity gene module GM_D, characterized by gene signatures connected to interleukin-17 (IL-17) producing T cells^13,26–28^, was exclusively enriched within c8, which contained neonatal as well as adult γδ T cells (**Fig. 2 and Supplementary Fig. 2b**). Finally, the major adult-derived γδ T cell clusters (c11 and c12) expressed genes related to NK cells (*KLRC1, KLRC1*), cytotoxicity (*FCGR3A, BATF, GNLY*), and granzymes (*GZMA, GZMB*) that confined them to the cytotoxic T lymphocyte (CTL) response gene module (GM_F) (**Fig. 2 and Supplementary Fig. 3**).

**Figure 2:**
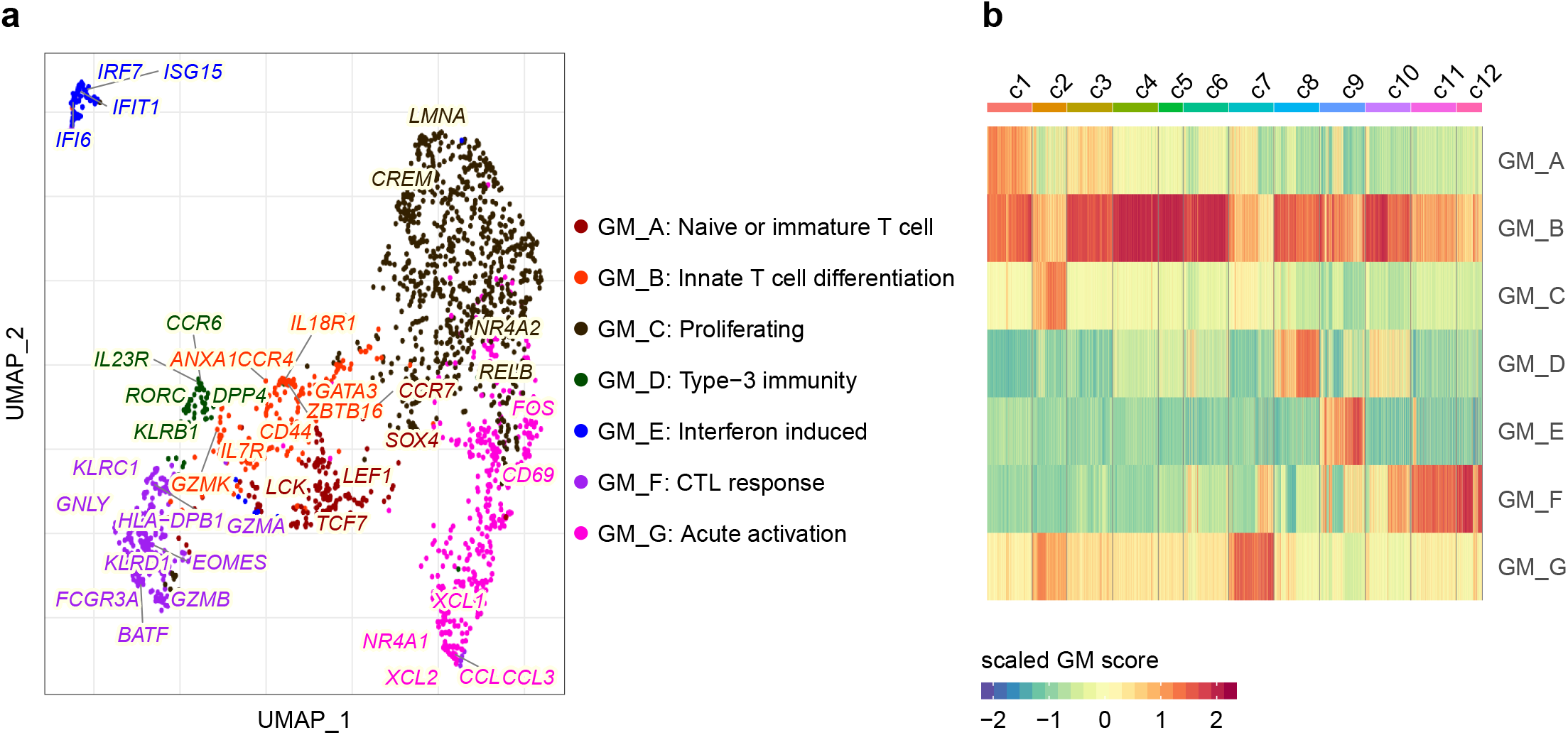
Identification of gene co-expression modules from single-cell transcriptomes. All 1,934 DEGs (logFC ≥ 0.25, adj.p ≤ 0.01, min-pct ≥ 10%, filtered ribosomal and mitochondrial genes) were subjected to unsupervised clustering based on the DEG’s average expression per cluster (c1-c12) and seven gene co-expression modules (GMs) were identified. Annotation of the seven GMs was supported by gene ontology enrichment analysis (Suppl. Fig3). (**a**) For data visualization, the seven GMs were enwrapped by UMAP and color-coded. Selected key genes were further labeled on UMAP. Each dot represents a single cell that originates from the respective donors in Fig1a-b. (**b**) The heat map depicts the modula scores of each cell. Each column reflects a single cell, and the color bars above represent c1-c12. Cell numbers were downsampled to max. 600 cells per cluster.

Altogether, scRNA-seq of > 20,000 human γδ T cells from cord blood and adult peripheral blood revealed a high functional heterogeneity with common and distinct cell transcriptional patterns among neonatal and adult γδ T cells. Detailed analysis of DEGs identified key core gene co-expression modules that defined a naïve phenotype of γδ T cells, their activation, differentiation to functional subsets, as well as type-3 and CTL effector functions.

### Expression of *TRD* clones clusters immature, Vδ1^+^, and Vδ2^+^ T cells

Human γδ T cells are typically classified according to their expressed Vδ-chain^3^. To link the single-cell transcriptome profiles described above to individual γδTCR clones, we next PCR-amplified and sequenced full-length V(D)J segments in a two-step enrichment process using gene-specific primers for the *TRDC* and *TRGC* genes as described in the methods section. Cellular barcodes allowed mapping of approximately 45 to 70% of single-cell transcriptomes to individual paired *TRG* and *TRD* clones, respectively (**Fig. 3a,b and Supplementary Fig. 4a**). Focusing on the more diverse *TRD* repertoires, the identified 12 clusters largely delineated the distribution of *TRDV* genes into three main sections. An immature section included neonatal γδ T cell clusters (c1 – c3) consisting of a heterogeneous mix of *TRDV1, TRDV2, TRDV3*, and a few other *TRDV* genes. The adult γδ T cell clusters c11 and c12 as well as the mixed neonatal/adult cluster c7 were mainly *TRDV1*^+^, while the *TRDV2*^+^ section was composed by cord blood-derived clusters c4 and c5, mixed clusters c6, c8, c9, and adult cluster c10 (**Fig. 3a,b**). As expected, those Vδ2^+^ clones preferentially paired with Vγ9 chains (**Supplementary Fig. 4b**). Next, we examined γδ T cell clonal expansions based on the distribution of individual paired CDR3γ and CDR3δ sequences (**Fig. 3c**) and corresponding Gini diversity indices (**Fig. 3d**). In general, adult Vδ1^+^ and Vδ2^+^ T cells showed a focused oligoclonal *TRD* repertoire (high Gini indices), while neonatal Vδ1^+^ and Vδ2^+^ *TRD* repertoires were rather polyclonal (lower Gini indices)^9,10,29^. Specifically, immature neonatal γδ T cells of the c1 – c3 section, which were regulated by gene modules of naïve or immature T cell (GM_A), showed no clonal expansions and Gini indices in c1 – c3 were close to zero (**Fig. 3c,d**). Strikingly, c11 cells, which mainly originated from the adult PB_donor2 (**Fig. 1b**), were largely constituted by one single *TRGV5/TRDV1* clone (**Supplementary Fig. 4a-c**), indicating that these cells accumulated due to antigen-specific clonal responses. However, we do not have any detailed information on the medical record of anonymous adult donors 1 and 2, so we can only speculate that these cells may have expanded in response to a viral infection such as CMV^9,10,30^. In sum, scTCR-seq introduced an additional layer of information and segregated the 12 γδ T cell clusters into immature, *TRDV1*^+^, and *TRDV2*^+^ sections that differed in their clonal compositions.

**Figure 3:**
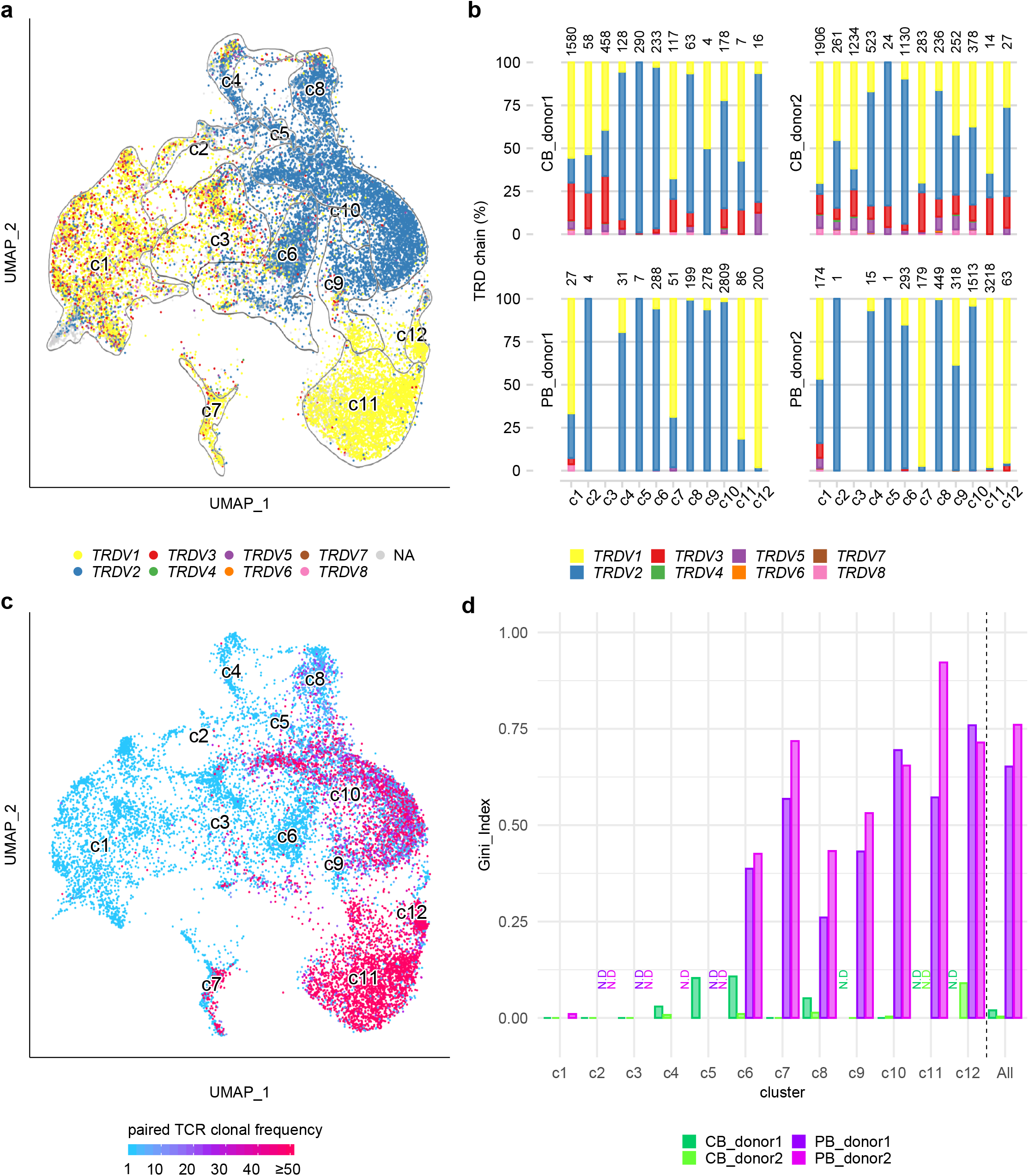
Single-cell TCR repertoire shows the correlation between transcriptome and TCR. Full-length γδTCR V(D)J segments were amplified from cDNA of scRNA-seq libraries for NGS analysis (n = 4). Via cellular barcodes single-cell γδTCR repertoires were mapped to matching single-cell transcriptomes of the respective four donors described in Fig1a. (**a**) UMAP of γδ T cells visualizes the *TRD* repertoires’ V gene usage (*TRDV*), while dots represent single-cells. Cells without productive *TRD* gene rearrangements are shown in grey. Contours outline cluster c1-c12 defined in Fig.1a. (**b**) Bar plots depict *TRDV* gene compositions in each cluster per donor. Cells without a productive *TRDV* gene were excluded. The total number of mapped *TRDV* genes per cluster are provided on top of each bar. (**c**) The frequency of each identified paired γδTCR clone was calculated and visualized within the UMAP of γδ T cells. Cells without a paired γδTCR were excluded. (**d**) Single-cell paired TCR repertoire diversities were estimated by Gini indices that range from 0 (completely polyclonal) to 1 (completely monoclonal). Bar plot shows Gini indices of each donor within the respective cluster (c1-c12) and total single-cell transcriptomes (all clusters per donor). Clusters containing less than 20 paired TCR were excluded from this analysis and defined as N.D (no data).

### Vδ1+ γδ T cells mature into PD-1hi and PD-1low cells

To understand the effector differentiation associated with clonal expansion of adult Vδ1 ^+^ T cells, we analyzed their transition from immature to effector Vδ1^+^ T cells at the transcriptional level. While Vδ1^+^ T cells of c1 were dominated by the gene module of naïve and immature T cells (GM_A), both neonatal and adult Vδ1^+^ T cells in c7 showed gene expression profiles of acute activation (GM_G). Adult effector Vδ1^+^ T cells (c11, c12) were controlled by the gene core module of CTL responses (GM_F) including *TBX21, EOMES*, and granzyme genes (**Fig. 2b and 4a**). Intriguingly, cells within the clonal cluster c11 expressed exhaustion signature genes such as *IKZF2, PDCD1, LAG3*, and *HLA-DRA*, whereas c12 was enriched for NK receptors, e.g. *KLRD1* and *KLRF1*, and cytotoxic genes, e.g. *FCGR3A* (CD16) and *TYROBP* (**Fig. 4b**). Based on these signatures, and supported by gene set enrichment analysis, adult Vδ1^+^ T cells of PD-1^hi^ c11 and PD-1^low^ c12 clusters can thus be classified into exhausted and active subsets, respectively (**Fig. 4c**).

**Figure 4:**
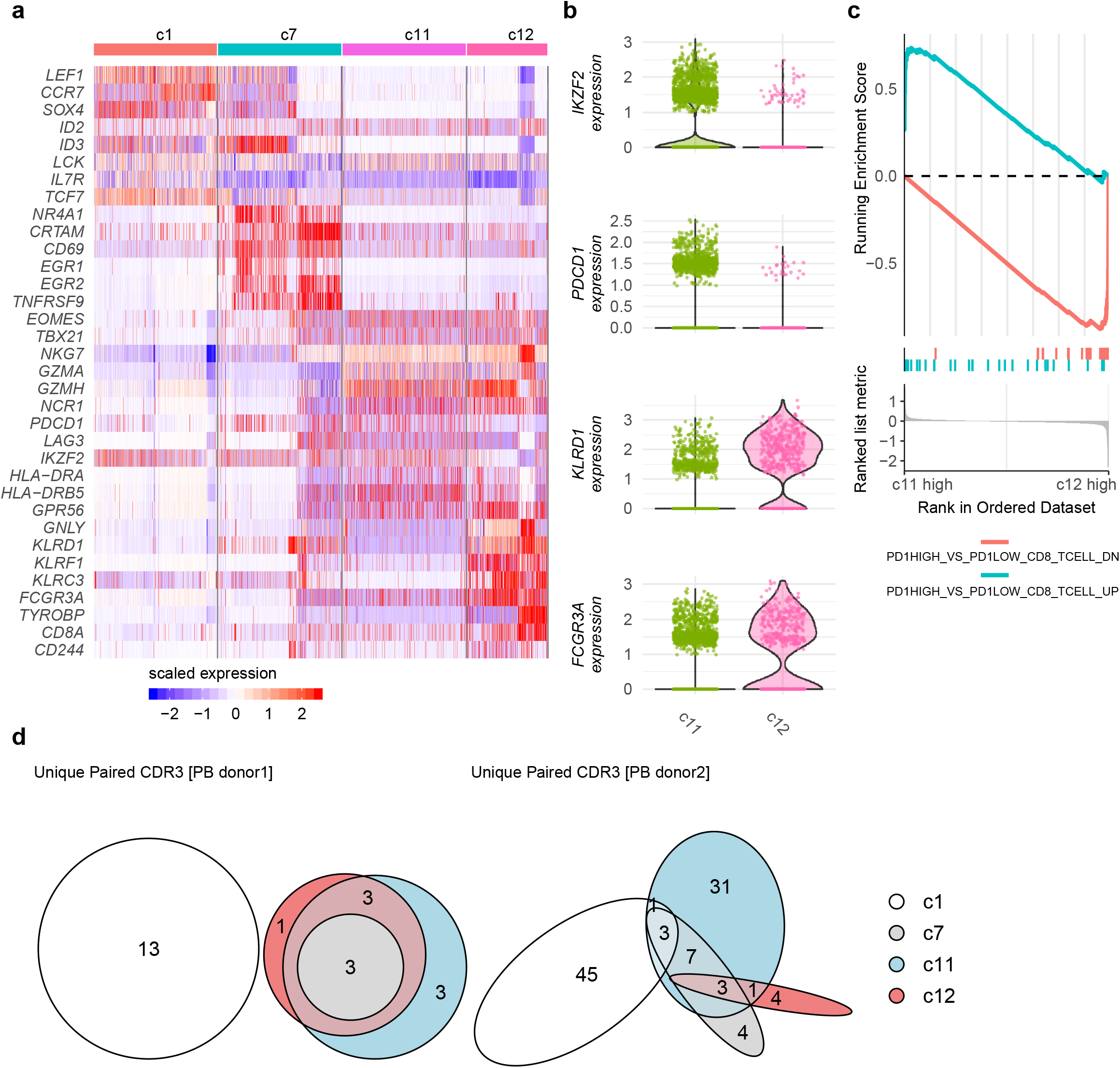
Vδ1^+^ γδCTLs mature into PD-1^hi^ and PD-1^low^ cells. Cell clusters with high *TRDV1* gene expression (from Fig. 3a; c1, c7, c11-c12) were further analyzed. (**a**) Heat map describes selected DEGs among cluster c1, c7 and c11-c12. (**b**) Expression levels of DEGs that are key markers of exhausted (*IKF2, PDCD1*) or activated (*KLRD1, FCGR3A*) T cells in c11-c12 were plotted by violin scatter plots. (**c**) Gene set enrichment analysis (GSEA) was performed on gene expression profiles of c11 versus c12 referring to the Immunological gene sets of Molecular Signatures Database (MSigDb; www.gsea-msigdb.org). The enriched terms were from an analysis of CD8^+^ T cells, GEO (GSE26495). Lower panel: Genes are arranged from being enriched in c11 to being enriched in c12 based on their logFC values. Middle panel: Each bar represents one hit gene from the enriched term on the gene list of c11 versus 12. Upper panel: Curves depict the running enrichment score of the selected enriched terms. (**d**) Paired TCR repertoire analysis of Vδ1 ^+^ T cells (c1, c7, c11 – c12) from PB donor_1 and PB donor_2. All paired TCR clones of the respective cluster were subjected to overlap analysis. Venn diagrams describe shared and non-shared TCR clones among clusters within PB donor_1 (left side) or PB donor_2 (right side). The size of each ellipse reflects the number of unique Vδ1^+^ clones in the respective cluster, further displayed in numbers.

Mapping of unique paired CDR3γ/CDR3δ sequences from the two adult donors to the Vδ1 ^+^ clusters revealed that the clones of c7, c11, and c12 were highly overlapping in each donor but distinct from immature cells in c1 (**Fig. 4d**). Altogether, these data support a trajectory of Vδ1^+^ T cells from an immature/naïve state in c1 over an acute activation in c7 to differentiate into fully activated Vδ1^+^ γδCTLs (c12, PD1^low^) or exhausted Vδ1^+^ γδCTLs (c11, PD1^hi^) in adult individuals.

### Identification of a distinct IL-17-committed Vδ2+ γδ T cell subset

Next, we investigated the heterogeneity of neonatal and adult Vδ2^+^ T cell subsets in the *TRDV2*-dominated clusters c4 – 6 and c8 – 10. Interestingly, when comparing the 3.764 individual *TRDV2* gene clones from this scTCR-seq analysis to a large number of *TRD* repertoires obtained from 80 independent donors^9,16^ by bulk TCR sequencing of Vγ9Vδ2^+^ T cells, we noticed that the majority of all Vδ2^+^ T cells displayed public *TRDV2* gene rearrangements, i.e. exactly the same *TRDV2-CDR3* sequences were shared between individual samples (**Fig 5a and Supplementary Fig. 5a**). This may have been a consequence of an expansion of public fetal-derived Vγ9Vδ2 T cell in the perinatal period^31,32^. In contrast, basically all *TRDV1* clones were private, i.e. not sharing their unique *TRDV1-CDR3* between individuals. Of note, Vδ2^+^ T cell clones of CB samples^16,33^ and of c8 showed high levels of public abundance (**Supplementary Fig. 5b**). In addition to their high level of shared *TRDV2* clones, these Vδ2^+^ T cells had *TRGV9/TRGJP* rearrangements (**Supplementary Fig. 5c**) and expressed two genes that guide lineage commitment of innate-like T cells including canonical Vγ9Vδ2^+^ T cells^15,34^, namely *ZBTB16* (encoding PLZF) and CD161-encoding *KLRB1* (**Fig. 5b**).

**Figure 5:**
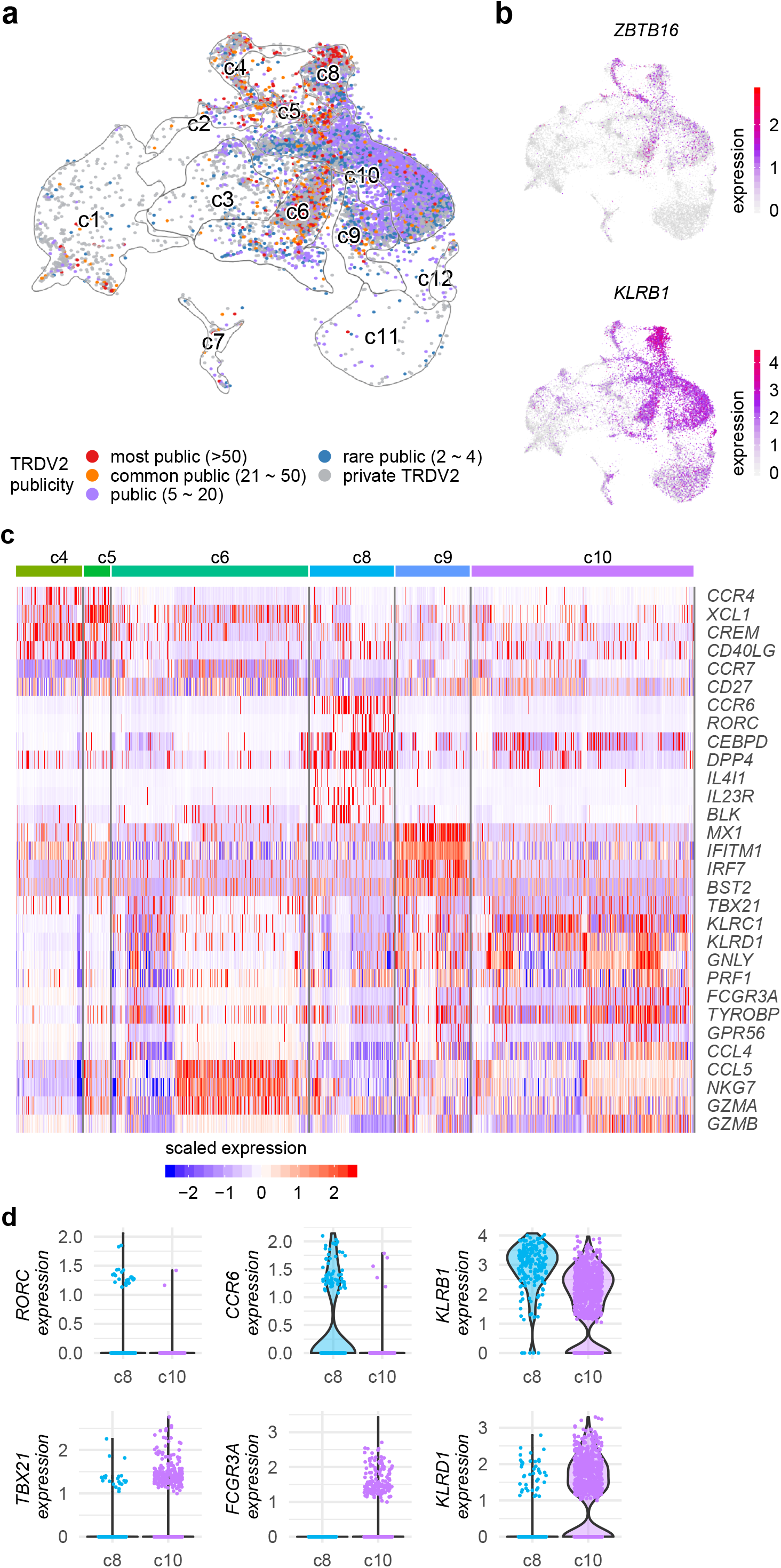
Vδ2^+^ γδ T cells are heterogeneous innate-like T cells. Combined scTCR-seq and scRNA-seq analysis focusing on neonatal and adult Vδ2^+^ T cell clusters defined in Fig. 3. (**a**) UMAP of Vδ2^+^ γδ T cells with delineation of respective clusters (from Fig. 1a) projects the publicity of *TRD* repertoires per cluster. Bulk *TRD* repertoires from independent study populations (ranging from infants to adults, n = 80) were compared to the four scTCR-seq libraries of this study and devoted to define a list of most public, common public, public, rare public and private Vδ2^+^ T cell clones by being present in the respective number of donors, indicated in brackets of the legend. (**b**) Visualization of *ZBTB16* (encoding PLZF) and *KLRB1* (encoding CD161) expression on γδ T cells (adopted from Figure S2c). (**c**) Cell clusters dominated by *TRDV2* expression (from Fig. 3a, c4 – c6, c8 – c10) were further investigated. Expression levels of selected DEGs were plotted by heat map. (**d**) Key DEGs of innate-like T cells (*KLRB1*), Th-17 (RORC, CCR6) and Th-1 (TBX21, FCGR3A, KLRD1) immunity are illustrated by violin scatter plots within c8 cells (cord blood and adult) and c10 cells (adult only).

Nevertheless, a number of other genes were differentially expressed among the Vδ2^+^ T cell clusters (**Fig. 5c,d**). The mostly CB-derived Vδ2^+^ T cells of c4 and c5 expressed *CCR4* and *CD40LG*, related to type-2 immunity^35^. In contrast, the mostly adult-derived Vδ2^+^ T cells in c9 expressed type-1 immunity signature genes such as *TBX21* and genes related to cytotoxicity, e.g. *FCGR3A* (CD16)*, PRF1*, and *GZMA*, suggesting differential roles of Vδ2^+^ T cells in the neonatal and adult immune system. However, clusters c6, c8, and c10 were composed of Vδ2^+^ T cells derived from both cord and adult blood donors. Of those, c6 cells displayed gene expression profiles (*CCR7, CD27*) of naïve T cells, and c10 cells were enriched in IFN-induced genes. Interestingly, c8 cells expressed type-3 immunity/Th17 signature genes including *CCR6, RORC, IL23R*, and *DPP4* (**Fig. 5c,d and Supplementary Fig. 5d**), thereby identifying c8 Vδ2^+^ T cells as IL17-committed γδ T cells (γδT17) that were present in neonates and adults.

To investigate the clonal relationship of naïve (c6), type-1 IFN-related (c9) or γδT17 (c8) Vδ2^+^ T cell subsets in adults with the γδT1 cells in c10, we calculated the overlap of paired CDR3γ/CDR3δ sequences. Strikingly, naïve (c6) and effector (c9, c10) Vδ2 T cells shared >50% to 100% of their clones in both adult samples while γδT17 cell (c8)-derived CDR3δ sequences were using distinct TCR clonotypes/sequences (**Fig. 6a**). Furthermore, none of the five most expanded Vγ9Vδ2^+^ *TRD* clones in c8 were found in c6, c9, and c10 clusters of the respective donors (**Fig. 6b**). Together, this establishes a close developmental relationship of IFN effector (c9) and adult γδT1 Vδ2 T cells (c10), while γδT17 cells (c8) show distinct/different yet public TCR repertoires. In combination with the type-3 immunity-related gene expression profile of c8 cells, this supports the hypothesis that human γδT17 cells may have a distinct early ontogenetic origin.

**Figure 6:**
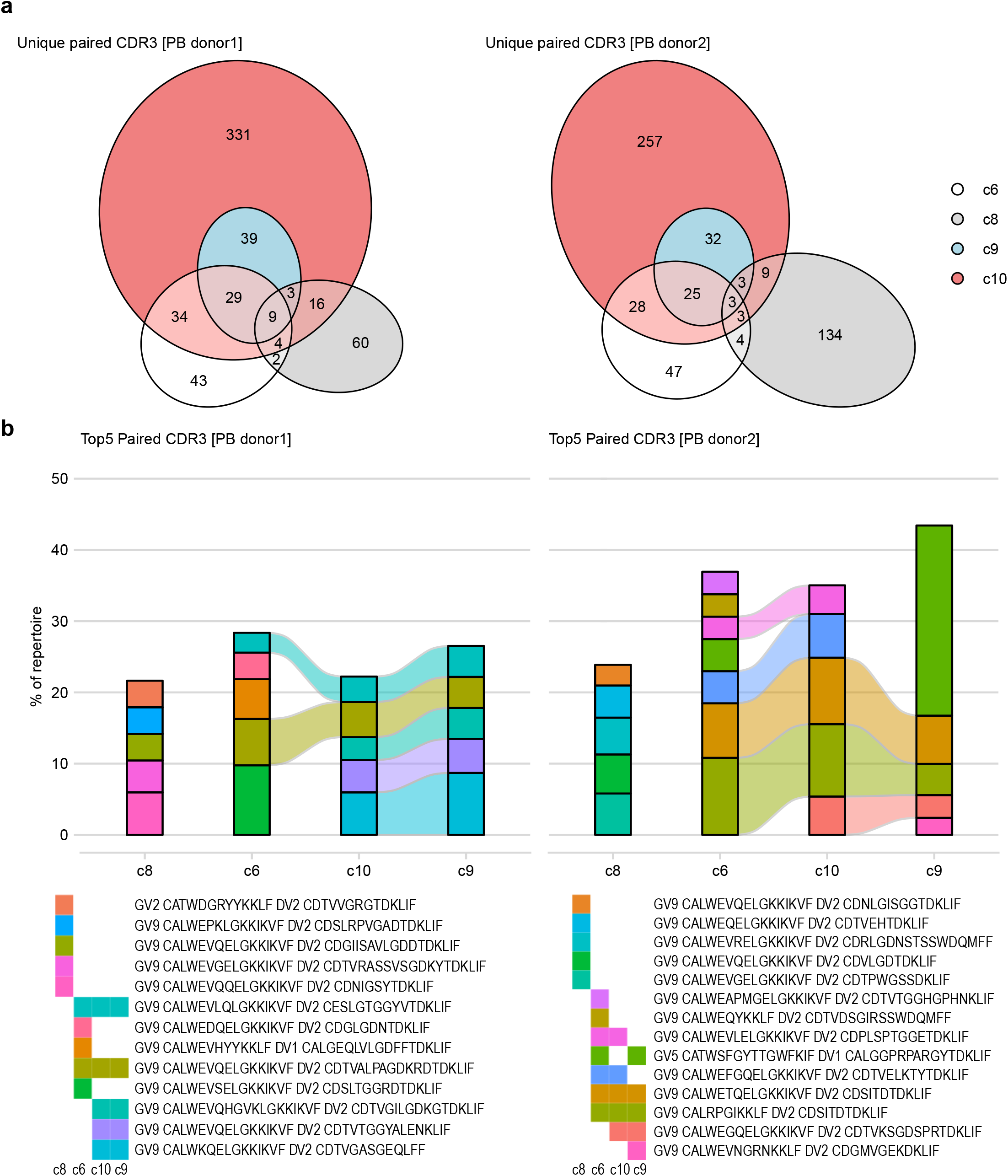
γδT17 cells show distinct TCR repertoire profiles. Paired TCR repertoires of Vδ2^+^ T clusters either being committed to a Th17-like (c8) or Th1-like (c6, c10, c9) response were investigated in adult PB_donor1 and PB_donor2. (**a**) Overlap analysis of paired TCR clones obtained from scTCR-seq was performed for PB_donor1 and PB_donor2. The amount of overlapping clones among clusters is visualized and described in venn diagrams, while the size of the ellipse reflects the number of unique TCR clones in each cluster. (**b**) The five most expanded paired TCR clones (Top-5) of each *TRDV2* dominated cluster (c8, c6, c10, c9) were selected for overlap analysis. The colored bands between two columns represent shared clones among clusters. The order of clusters was adjusted for visualizing. Results are described for PB_donor1 on the left side and for PB_donor2 on the right side. (**a,b**) Cluster c4 and c5 were excluded from analysis as few cells originated from adult donors.

### Human γδT17 cells exclusively originate from the earliest embryonic thymocytes

Murine γδT17 cells emerge during a defined window of embryonic development and are prewired to produce IL-17^12,28^. To test whether human γδT17 cells arise similarly during early human ontogeny, we took advantage of a recent scRNA-seq study that resolved human thymus organogenesis in the embryonic weeks 8 to 10 (Ewk8 - 10)^36^. Re-analyzing this valuable dataset (GEO:GSE133341), we identified four thymic γδ T cell clusters (gdT_1 to 4) according to *CD3D* and *TCR* gene expression (**Fig. 7a and Supplementary Fig. 6**). Most strikingly, two smaller γδ T cell clusters (labeled gdT_3 and gdT_4) exclusively originated from Ewk8 and Ewk9 thymi, while the two larger γδ T cells cluster (labeled gdT_1 and gdT_2) mainly arose from Ewk10 thymocytes (**Fig. 7b**). Interestingly, applying the gene modules identified above (**Fig. 2**), the majority of Ewk10 γδ T cells (gdT_1 & 2) as well as the αβ T cell clusters were distinguished by gene modules of naïve and immature T cells (GM_A) (**Fig. 7c**). In contrast, γδ T cells enriched in Ewk8 and Ewk9 (gdT_3 and 4), showed strong expression of the innate T cell differentiation gene module (GM_B). Moreover, the Ewk8 and Ewk9 clusters were separated by CTL response (γδT_4) and type-3 immunity (gdT_3) gene modules (**Fig. 7c**). The heat map in **Fig. 7d** highlights the differential expression of cytotoxicity-related genes (e.g. *KLRD1, PLAC8, NKG7, TYROBP, GZMA*) on cluster gdT_4 and of γδT17 cell signature genes (e.g. *RORC, CCR6, IL23R*) on cluster gdT_3. Both subsets expressed *ZBTB16* (encoding PLZF) and *KLRB1* (encoding CD161), two genes previously described on fetal-blood Vγ9Vδ2 T cells^15^. Expression of *CD44* and *CD69* suggests that these two γδ T cell subsets are already activated in the embryonic thymus (**Fig. 7d**). In addition, re-analysis of embryonic thymocytes from the human cell atlas^37^ also revealed a small group of γδ T cells from embryonic week 7 that expressed GM_B: Innate T cell differentiation and GM_D: Type-3 immunity (not shown). However, the low number of γδ T cells in that dataset made it less evident than re-analysis of the Zeng et al. resource^36^.

**Figure 7:**
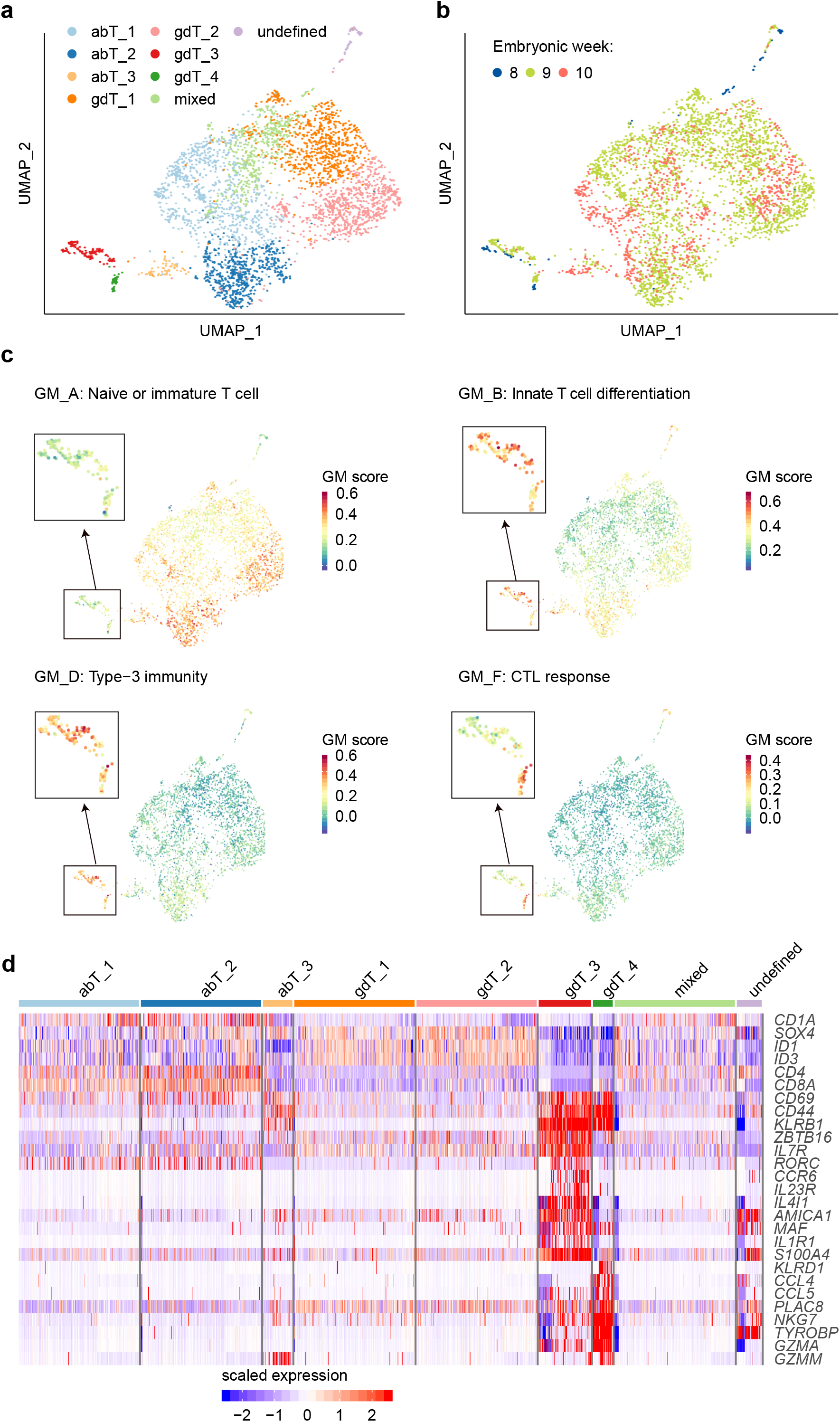
γδ T cells acquire effector phenotypes during the early fetal thymic development. Single-cell transcriptome dataset of thymi from embryonic week 8 to 10 was acquired from Gene Expression Omnibus (Zeng et al., 2019; GEO: GSE133341). T cells (defined by *CD3D* expression in Fig.S6a) were subjected to this re-analysis. (**a**) UMAP illustrates unsupervised clustering results of fetal-thymus T cells. Names of clusters were determined by the expression of TCR genes (Figure S6b). (**b**) UMAP describes the origin in embryonic weeks of fetal-thymus T cells via respective color codes. (**c**) Aggregated expression values of GMs (defined in Fig. 2) were projected on the UMAP of fetal-thymus T cells. Cluster gdT_3 and gdT_4 are magnified in the black box for clearer visualization. (**d**) Representative key DEGs among fetal-thymus T cells are presented in heat map. Cells were downsampled to max. 200 cells per cluster.

In summary, human γδT17 cells were exclusively present in the human thymus at embryonic week 8 and 9, but not week 10. Their transcriptional patterns point towards a functional precommitment for IL-17 production during early thymic development that is conserved and persists in adult peripheral blood γδT17 cells.

### Delineating Vδ1^+^ and Vδ2^+^ effector subsets via flow cytometry

The combination of scRNA-seq and scTCR-seq identified a number of distinct phenotypes of human γδ T cells, including γδT17 cells, in cord blood and adult peripheral blood. To validate and apply these results, we selected nine differentially expressed indicator surface markers that represented the phenotypes and gene modules of γδ T cells for validation by flow cytometry (**Supplementary Fig. 7**). In addition to antibodies specific for Vδ1^+^, Vδ2^+^, and pan-γδTCR, the panel included nine antibodies directed against CCR7 and CD127 (*IL7R*) for central memory and naïve phenotypes, CCR4 to identify Th2-like Vδ2^+^ T cells, CD161 (*KLRB1*), CCR6, and CD26 (*DPP4*) as markers for γδT17 cells ^38,39^, CD94 (*KLRD1*) and CD16 (*FCGR3A*) representing CTL activity, and PD-1 (*PDCD1*) indicating exhaustion. With this simplified approach, we could successfully reproduce the presence of γδT17 and other γδ T cell subsets in independent cord and adult blood donors. Based on the combined flow cytometric analysis of five cord blood and six adult blood samples by UMAP and unsupervised clustering, we identified ten distinct clusters (named FACS1-10) that largely corresponded with the twelve clusters identified by scRNA-seq (**Fig. 8a,b and Supplementary Fig. 8a-c**). Notably, although dimensional reduction was performed based on only nine selected surface markers, the clustering obtained from this analysis segregated sections of immature (mixed), Vδ1^+^ and Vδ2^+^ T cells (**Fig. 8c**) similarly to total scRNA-seq analyses (**Fig. 3a**). Also, γδ T cells allocated to distinct clusters depending on their origin from neonatal or adult donors (**Fig. 8d and Supplementary Fig. 8a**). A prominent exception to this rule was cluster FACS6, representing CCR6^+^CD26^+^CD161^+^Vδ2^+^ γδT17 cells. To address whether these cells comprised in cluster c8/FACS6 could be generated from hematopoietic stem cells in a postnatal thymus, we applied our FACS panel to analyze the peripheral blood γδ T cells of an adult *IL2RG*-deficient severe combined immunodeficiency (SCID) patient who was transplanted with *IL2RG*-sufficient T cell-depleted bone marrow at the age of one (**Supplementary Fig. 8d**). In line with previous reports^40^, frequencies of Vγ9^+^CCR6^+^ T cell frequencies were highly heterogeneous in healthy control donors, however basically undetectable in the reconstituted SCID patient.

**Figure 8:**
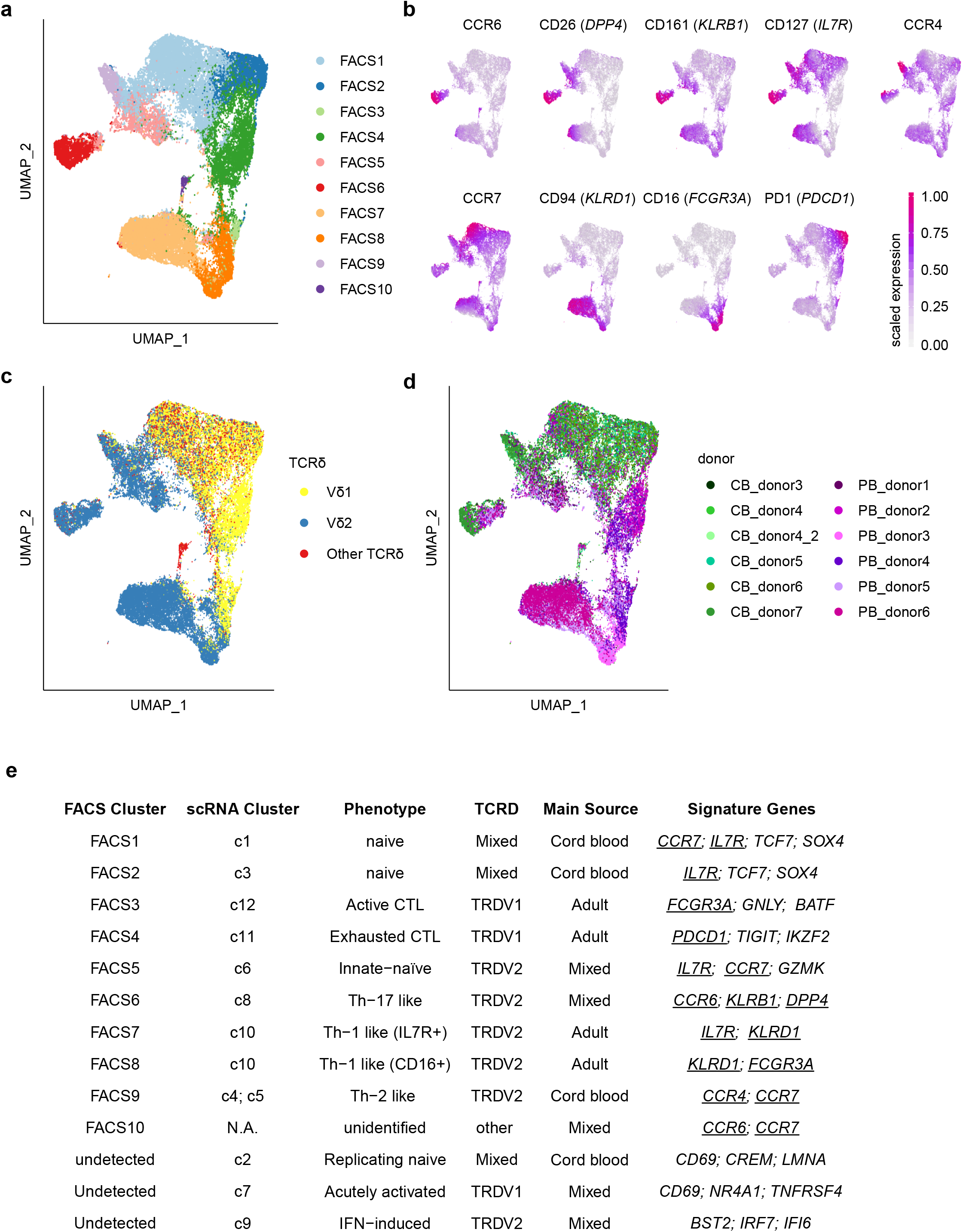
Flow cytometry analysis delineates Vδ1^+^ and Vδ2^+^ naïve and effector γδ T cell subsets. γδ T cells from five cord blood and six adult peripheral blood donors were subjected to multicolor FACS analysis. CB_donor4_2 was an independent replicate of CB_donor4 that serves as an internal control for batch effects, while PB_donor1 and PB_donor2 were identical to adult donors of scRNA-seq data. Neonatal and adult γδ T cell were stained by pan-γδTCR, Vδ1 and Vδ2 antibodies and nine antibodies for key surface markers to distinguish different phenotypes were included (**a**) FACS data of γδ T cells was subject to unsupervised clustering that was based on those nine key surface markers and enwrapped by UMAP. Ten FACS clusters were identified. Each point represents one single-cell and is color-coded by FACS cluster. (**b**) Expression values of the nine lineage markers were scaled to a range between 0 and 1 and projected on UMAP. (**c,d**) Individual cells are color-coded by (c) corresponding TCRδ chains or (d) donor ID. Gating of Vδ1^+^ and Vδ2^+^ T cells is shown in **Supplementary Fig. 8b**. (**e**) Overview of the corresponding clusters from multicolor FACS analysis and scRNA-seq, phenotypes, sources, TCRδ usage and corresponding signature genes are summarized in the table. The nine lineage markers selected for FACS validation are underscored.

The phenotype, composition, key markers and interrelatedness of γδ T cell clusters defined by flow cytometry versus scRNA-seq is summarized in **Fig. 8e**. In brief, the flow cytometry panel comprising three γδTCR and nine surface molecule markers not only validated the prior scRNA-seq data, but provides a simple strategy to monitor the diverse γδ T cell subsets identified in neonatal and adult blood.

## DISCUSSION

In this work, scRNA-seq of sorted adult and neonatal γδ T cells allowed us to establish a comprehensive map of γδ T cell phenotypes and to associate them to paired γδTCR sequences. In essence, functional diversity of human γδ T cells, as monitored by DEGs, largely correlated with the type of γδTCRs expressed in each cell. While polyclonal neonatal cells dominated the more “naïve” first half of clusters (c1 to c6), characterized by expression of genes involved in innate T cell differentiation, the other half of clusters (c7 to c12) showed distinct patterns of T cell activation, proliferation, and lineage-specific differentiation. Furthermore, we extracted seven GMs^23,24^ that explained the naïve, proliferating, type-3 immunity, cytotoxic response, IFN-induced response, NKT differentiation, and acute activation phenotypes in human γδ T cells. Thereby, compositions of naïve, IFN-induced, and cytotoxic GMs resembled those identified from human αβ T cells^23^. Most importantly, our GMs allowed us to assess and interpret the phenotypes of embryonic γδ thymocytes from Zeng et al^36^, and we suggest that these GMs will serve as a universal tool to understand the transcriptional programming of human γδ T cells. Furthermore, we provide a FACS panel that, in addition to TCR-specific mAbs, requires only nine surface makers to largely reproduce the phenotypic map obtained by scRNA-seq.

A previous single cell analysis study^21^ identified γδ T cells in sets of human circulating and tumor-infiltrating cells. They suggested that Vδ1^+^ and Vδ2^+^ subsets form close yet distinct subclusters that similarly undergo a “cytotoxic maturation”, i.e. a gradual acquisition of CTL capacity. Our single cell analysis of large numbers of sorted γδ T cells and simultaneous analysis of their paired *TRG/TRD* repertoires now adds several levels of resolution to the understanding of human γδ T cell subsets.

Firstly, comparison of cord and adult blood samples was consistent with findings that neonatal γδ T cells are more immature and polyclonal^16,33,41^, whereas adult γδ T cells are more activated and show restricted clonality^9,10,42^. At the same time, cord blood and early fetal thymus contained innate γδT17 and γδT1 effector populations that presumably persisted into adulthood, sharing some features with Lin28b-dependent effector γδ T cells, recently discovered in week 17-19 fetal thymus^20^.

Regarding the distribution of clonotypes, cord (and adult) blood-derived γδ T cells in the immature subsets generally displayed a diverse Vδ1-dominated repertoire, with a high frequency of TCRs containing *TRDV3* and other non-*TRDV1* or *TRDV2* rearrangements. The high expression of *SOX4, ID2*, and *ID3* thereby suggests that these cells recently emigrated from thymus^10,18,41^. From the naïve state, we found a stepwise activation trajectory of Vδ1^+^ T cells that goes hand-in-hand with clonal expansion and expression of genes associated with CTL response, but also acquisition of exhaustion markers like PD-1 similar to CD8^+^ CTLs after intensive antigen-exposure^43^. Together with the notion that repertoire focusing and proportions of Vδ1^+^ CTLs vary drastically among adults^10^, this fits well with the hypothesis that naive Vδ1^+^ T cells differentiate into Vδ1^+^ CTLs and clonally expand after antigen-specific TCR-dependent activation.

In contrast, canonic Vγ9Vδ2^+^ T cells rather grouped with each other but separated into three effector types with strong parallels to lineage-specific differentiation of other PLZF^+^ innate lymphocytes such as NKT-1, NKT-2, and NKT-17 cells^44^. As an important disclaimer, certainly not all Vδ2-chains must pair with Vγ9-chains and *vice versa*^3,9,20,29^. However, besides the expression of PLZF, an abundance of public TCR clones, polyclonal expansion after phosphoantigen stimulation, and the lack of an exhausted phenotype suggest an innate rather than adaptive phenotype of Vγ9Vδ2^+^ cells^16^.

While Vγ9Vδ2^+^ cells with type-2 immunity-related markers including *CCR4* and *CD40LG* were rather restricted to the neonatal samples, we observed committed γδT17 and γδT1 cells already in the earliest wave of thymus organogenesis, as well as in cord blood and adult blood samples. Notably, this is consistent with the hypothesis that human innate Vγ9Vδ2^+^ effector γδ T cells develop in early fetal waves and persist into adulthood, as it has been described for mouse γδT17 and γδT1 cells^12,45^. So why was this not described before? Timing might be critical for their observation, since Vγ9Vδ2^+^ cells are certainly present very early in T cell ontogeny ^14,15^ but hard to detect in later fetal thymus (week 17-19)^20^ and absent in postnatal thymus^19^. In conclusion, the earliest innate γδT17 and γδT1 cells probably leave the human fetal thymus and home to distant tissues already after gestational week 9, in line with our detection of these cells in thymi of week 8 and 9, but not week 10. Such a strict fetal origin of effector cells is literally a feature of innateness and is in parallel to the ontogeny of IL-17-producing lymphoid tissue-inducer cells that seed and persist in secondary lymphoid tissues^46^. Likewise, PLZF expression maps the early stages of ILC1 lineage development^47^.

Furthermore, there is currently no uniform definition of human γδT17 cells. Several reports identified that subsets of Vδ2^+^ cells expressing IL-23R^48^, CD161 ^38,49^ or CCR6^38,40,50,51^ are the ones prone to IL-17 production. However, these reports concluded that peripheral differentiation of Vγ9Vδ2^+^ T cells into IL-17 producers required a highly inflammatory milieu. Here we define human type-3 Vγ9Vδ2^+^ T cells as γδT17 cells by expression of canonical Vγ9Vδ2^+^ TCR and the GM type-3 immunity comprising *IL23R, RORC*, and *BLK*, and by FACS as CCR6^+^, DPP4 (CD26)^+^, KLRB1 (CD161)^+^, and negative for KLRD1 (CD94) and FCγR3a (CD16). However, in sum it emerges that CCR6, like in mice^52^, is the most stringent single marker for human γδT17 cells.

A very recent report^53^ designated CD26^+^CD94^-^ Vδ2^+^ cells as a distinct population with a certain overlap to previously described CD28^+^CD16^-^ Vδ2^+^ cells^40^. Notably, both studies concluded that the earliest neonatal Vδ2^+^ cells are enriched in cells with an CD26^+^CD28^+^CCR6^+^ phenotype, but would later transition into a more mature profile of Vδ2^+^ cells^40,53^. However, from our scRNA-seq and FACS data it is evident that CD26^+^CD94^-^ Vδ2^+^ cells are actually a heterogeneous population comprising CCR6^+^ and CCR6^-^ cells.

Consistent with the finding of Ryan et al.^40^, we found that there are high levels of phenotypic variability between donors, with some profiles dominated by CCR6^+^ Vδ2^+^ cells and others that displayed only very few CCR6^+^ Vδ2^+^ cells, i.e. around 1%. However, we would speculate that these profiles are already established very early in ontogeny, perhaps in a neonatal window of opportunity, and persist and influence immune responses during an individual’s entire life. Only the analysis of larger long-term cohorts monitoring individuals from infancy to adulthood will be able to address this issue.

Notwithstanding, we propose the obvious, namely that human CCR6^+^ γδT17, similarly to mouse CCR6^+^ γδT17 cells^12^, are preferentially or even exclusively generated early before birth. This hypothesis is supported by i) the finding that an adult *IL2RG*-deficient SCID patient who was transplanted with *IL2RG*-sufficient T cell-depleted bone marrow at the age of one was lacking CCR6^+^ Vγ9^+^ T cells, ii) the direct identification of these cells in published dataset^36^ of fetal thymus Ewk8-9, but not Ewk10, and iii) the presence of the type-3 immunity-biased cluster c8 cells shared in samples from cord and adult blood, and iv) paired TCR analyses revealing that c8 clusters showed unique TCR profiles different to the other Vγ9Vδ2^+^ cell-dominated clusters (c6, c9, c10) with more overlapping TCR repertoires. Nevertheless, the physiological functions of human γδT17 cells remain to be established.

Taken together, we present a comprehensive map of functional human γδ T cell subsets and provide moderately complex FACS panel that can be easily implemented to reproduce this classification. This should contribute to decipher the factors that drive the individual TCR and functional subset composition of individuals early in life and to understand whether these parameters would influence or predict how individuals will cope with neoplastic and infectious challenges.

## Supporting information

all supplemental figures

## Acknowledgements

The work was supported by the Deutsche Forschungsgemeinschaft (DFG, German Research Foundation) under Germany’s Excellence Strategy – EXC 2155 “RESIST” – Project ID 39087428 to R.F., S.R. and I.P.; SFB900 – Project ID 158989968 to R.F., C.Könecke, S.R. and I.P. and DFG-funded research group FOR2799 to S.R. (Project ID RA3077/1-1) and I.P. (Project ID PR727/11-1). We would like to thank Matthias Ballmeier of the MHH cell sorting facility and the Genomics Core Unit of Hannover Medical School.

## Author contributions

L.T., A.F., S.R. and I.P. designed the study and experiments; L.T. analyzed data; A.F. contributed in data analysis; A.F., I.O., A.B. and S.R. conducted experiments; M.J. provided bioinformatics analysis tools; C.S.-F., C.Könecke and C.v.K. recruited and coordinated study participants; R.F., C. Krebs and U.P. provided technical support and intellectual input and L.T., A.F., S.R. and I.P. wrote the manuscript.

