## Supplementary material for "A fetal wave of human type-3 γδ T cells with restricted TCR diversity persists into adulthood": all supplemental figures

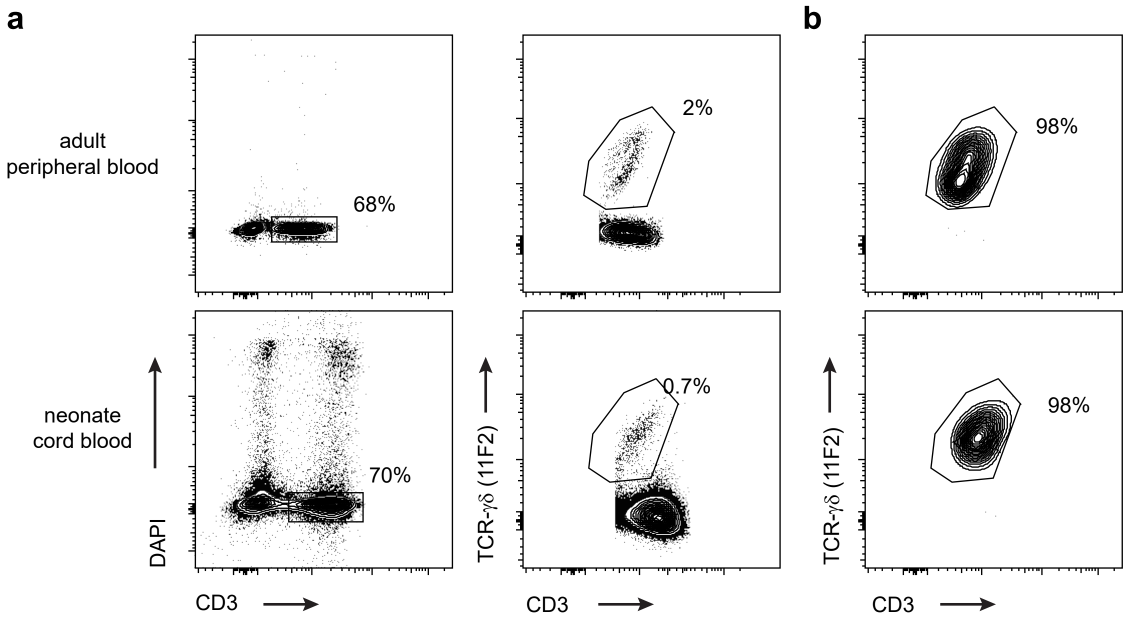


**Figure S1: Isolation of human γδ T cells for single cell NGS.**

(**a**) Peripheral blood from healthy adult donors and cord blood samples were collected and mononuclear cells were isolated by a density gradient. Representative FACS analysis to isolate living $CD3^{+}TCR\gamma\delta^{+}$ cells for scRNA-seq and scTCR-seq library generation. (**b**) Sorted γδ T cells were re-analyzed to validate cell purity.

Figure S2

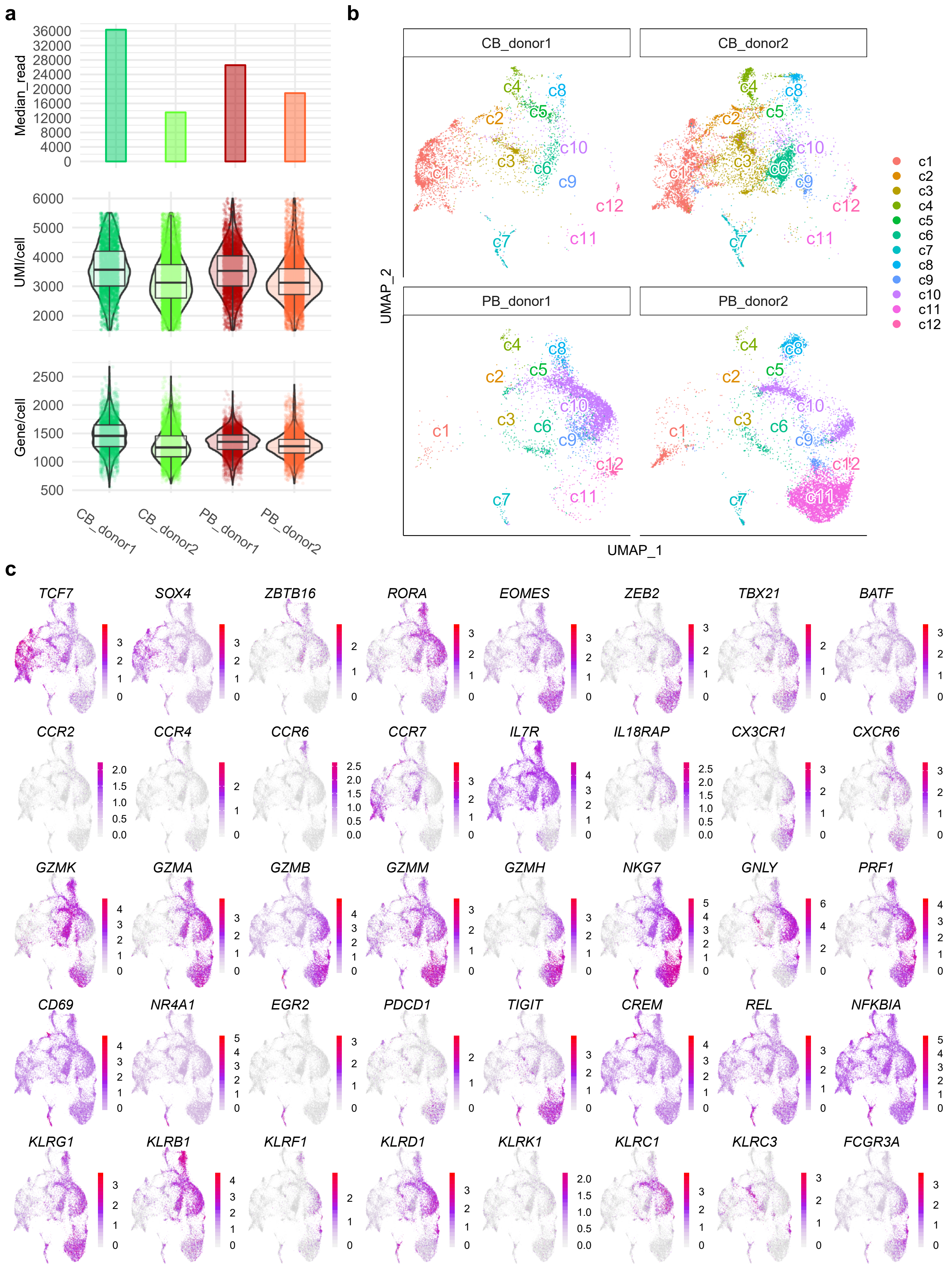
a
Median_read
UMI/cell
Gene/cell
CB_donor1
CB_donor2
PB_donor1
PB_donor2
b
CB_donor1
CB_donor2
PB_donor1
PB_donor2
UMAP_2
UMAP_1
c1
c2
c3
c4
c5
c6
c7
c8
c9
c10
c11
c12
c
TCF7
SOX4
ZBTB16
RORA
EOMES
ZEB2
TBX21
BATF
CCR2
CCR4
CCR6
CCR7
IL7R
IL18RAP
CX3CR1
CXCR6
GZMK
GZMA
GZMB
GZMM
GZMH
NKG7
GNLY
PRF1
CD69
NR4A1
EGR2
PDCD1
TIGIT
CREM
REL
NFKBIA
KLRG1
KLRB1
KLRF1
KLRD1
KLRK1
KLRC1
KLRC3
FCGR3A

**Figure S2: Quality-control and overview of scRNA-seq data.**

(**a**) Quality control of the four scRNA-seq library via obtained total reads as box plots, Unique Molecular Identifier (UMI) and detected gene number per cell within each donor as violin scatter and box-whisker plot. The three horizontal lines of the box-whisker plot represent the higher quartile, median, and lower quartile, respectively. The whiskers stretch from each quartile to the maximum or minimum. (**b**) UMAP of γδ T cells from **Figure 1a** is split by donor ID. (**c**) Expression levels of selected genes are projected on UMAP. Genes were ordered from transcriptional factors, chemical and interleukin receptors in the upper panel to cytotoxic proteins, T cell activation-related genes and NK cell markers in the lower panel.

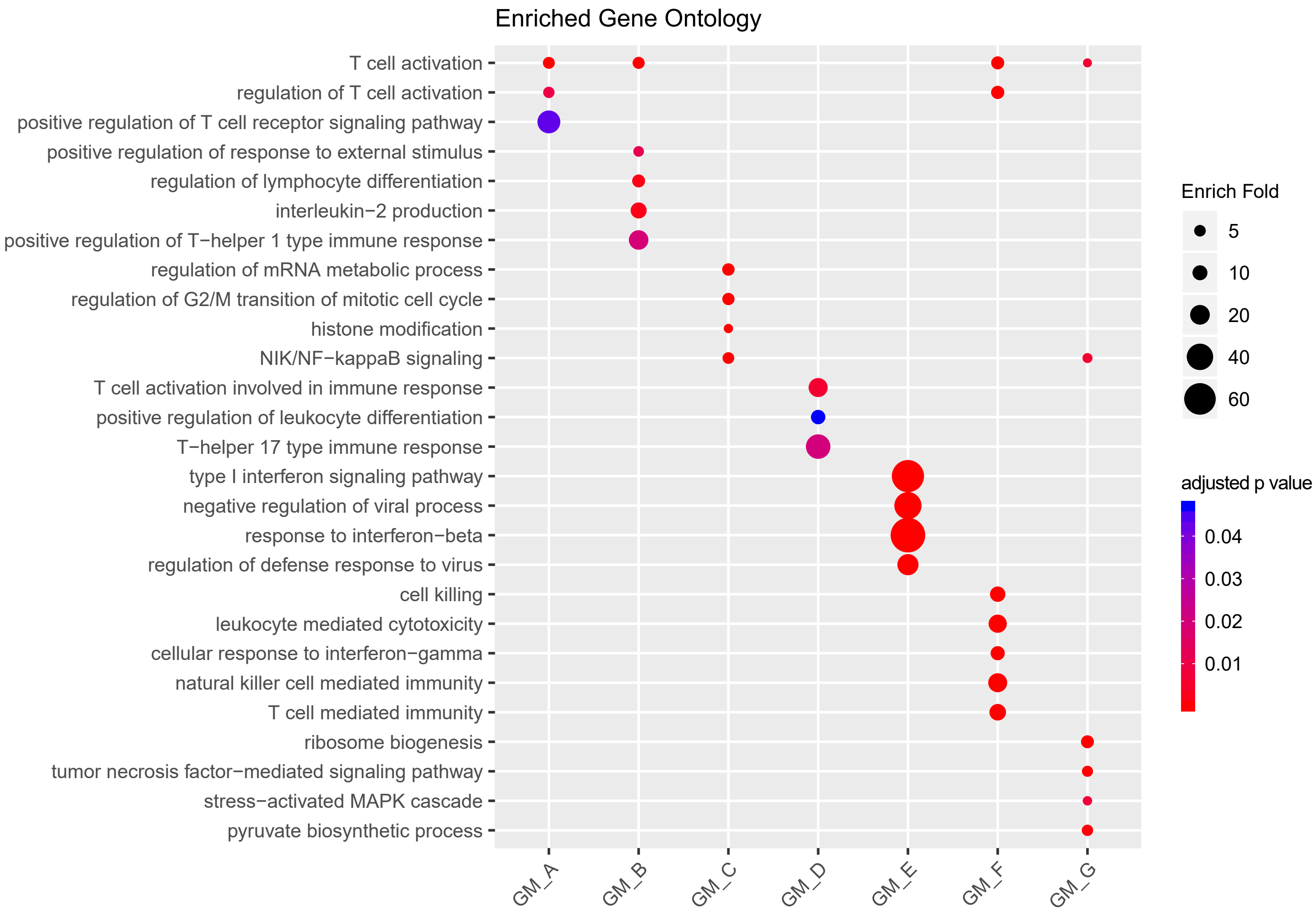


**Figure S3: Gene Ontology enrichment analysis on GMs.**
Genes in each gene co-expression modules (GM) were subjected to Gene Ontology (GO; biological process) enrichment analysis. Over Representation Analysis was used to identify the enriched GO terms. P-values were adjusted by Benjamini-Hochberg method. Selected enriched GO terms are presented on dotplot. The dots are colored by adjusted p-value and sized by enrich-fold respectively.

Figure S4

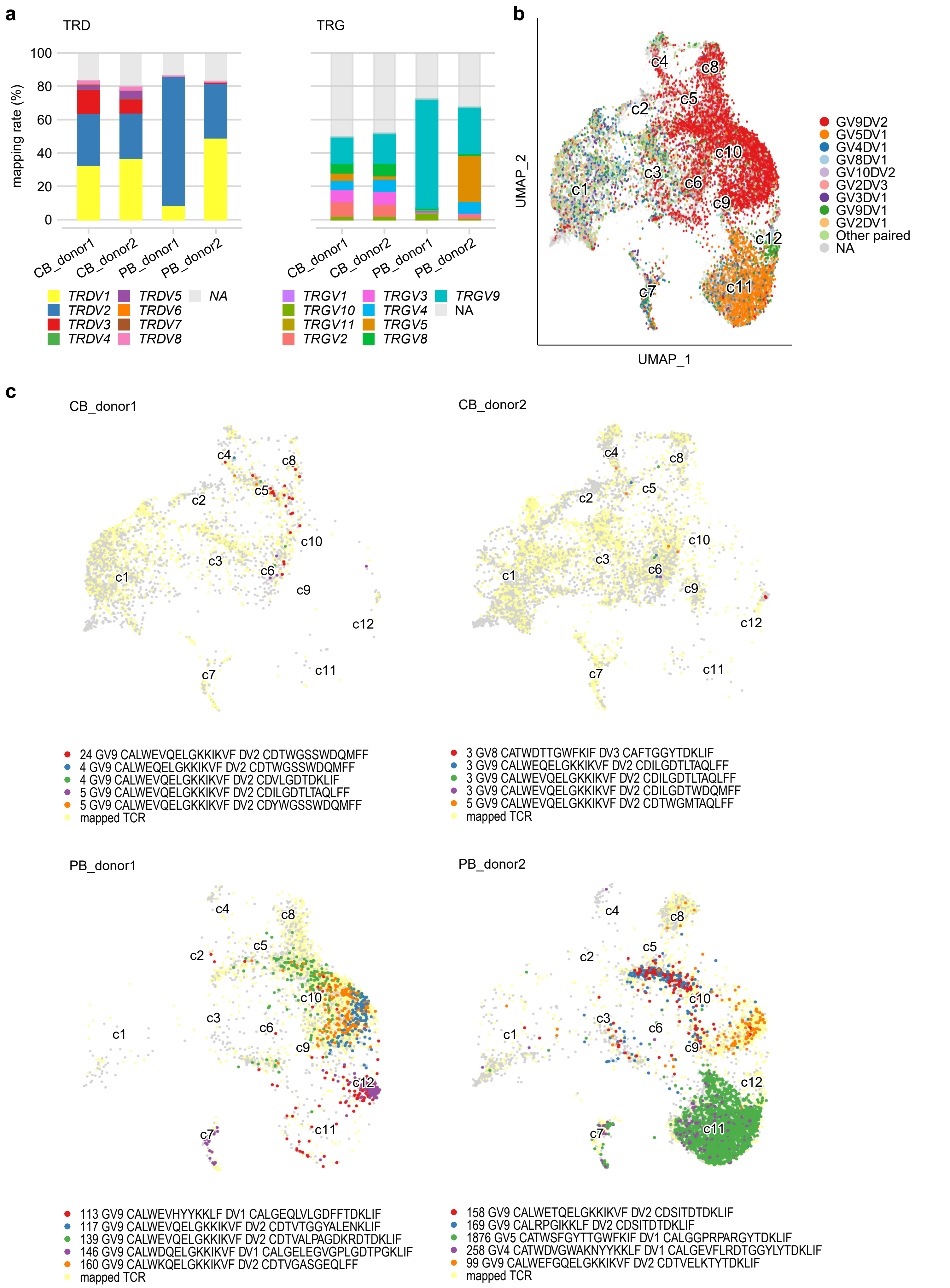
a
TRD
TRG
mapping rate (%)
100
80
60
40
20
0
CB_donor1
CB_donor2
PB_donor1
PB_donor2
TRDV1
TRDV2
TRDV3
TRDV4
TRDV5
TRDV6
TRDV7
TRDV8
NA
TRGV1
TRGV10
TRGV11
TRGV2
TRGV3
TRGV4
TRGV5
TRGV8
TRGV9
NA
b
UMAP_2
UMAP_1
c1
c2
c3
c4
c5
c6
c7
c8
c9
c10
c11
c12
GV9DV2
GV5DV1
GV4DV1
GV8DV1
GV10DV2
GV2DV3
GV3DV1
GV9DV1
GV2DV1
Other paired
NA
c
CB_donor1
24 GV9 CALWEVQELGKKIKVF DV2 CDTWGSSWDQMFF
4 GV9 CALWEQELGKKIKVF DV2 CDTWGSSWDQMFF
4 GV9 CALWEVQELGKKIKVF DV2 CDVLGDTDKLIF
5 GV9 CALWEVQELGKKIKVF DV2 CDILGDTLTAQLFF
5 GV9 CALWEVQELGKKIKVF DV2 CDYWGSSWDQMFF
mapped TCR
CB_donor2
3 GV8 CATWDTTGWFKIF DV3 CAFTGGYTDKLIF
3 GV9 CALWEQELGKKIKVF DV2 CDILGDTLTAQLFF
3 GV9 CALWEVQELGKKIKVF DV2 CDILGDTLTAQLFF
3 GV9 CALWEVQELGKKIKVF DV2 CDILGDTWDQMFF
5 GV9 CALWEVQELGKKIKVF DV2 CDTWGMTAQLFF
mapped TCR
PB_donor1
113 GV9 CALWEVHYYKKLF DV1 CALGEQLVLGDFFTDKLIF
117 GV9 CALWEVQELGKKIKVF DV2 CDTVTGGYALENKLIF
139 GV9 CALWEVQELGKKIKVF DV2 CDTVALPAGDKRDTDKLIF
146 GV9 CALWDQELGKKIKVF DV1 CALGELEGVGPLGDTPGKLIF
160 GV9 CALWKQELGKKIKVF DV2 CDTVGASGEQLFF
mapped TCR
PB_donor2
158 GV9 CALWETQELGKKIKVF DV2 CDSITDTDKLIF
169 GV9 CALRPGIKKLF DV2 CDSITDTDKLIF
1876 GV5 CATWSFGYTTGWFKIF DV1 CALGGPRPARGYTDKLIF
258 GV4 CATWDVGWAKNYYKKLF DV1 CALGEVFLRDTGGYLYTDKLIF
99 GV9 CALWEFGQELGKKIKVF DV2 CDTVELKTYTDKLIF
mapped TCR

**Figure S4: Overview of scTCR-seq data.**

(**a**) Proportions of cells of each donor that could be connected to a productive *TRD* (left) or *TRG* (right) clone according to the V-gene usage. (**b**) UMAP of γδ T cells colored by TRD & TRG combination of paired TCRs. Top-9 most abundant combinations were shown in different color code. NA: cells without a paired TCR. **(c)** Separate analysis of TCR repertoires of all four donors. All mapped paired TCR (yellow) are projected on the UMAP. The five most expanded (Top-5) paired TCR clones within each donor are visualized by different colors. The number of cells carrying the corresponding CDR3 sequences are specified in the legend.

Figure S5

**a**

| publicity | TRDV2 clones | Other TRD clones |
| --- | --- | --- |
| most public (>50) | 40 | NA |
| common public (21 ~ 50) | 201 | NA |
| public (5 ~ 20) | 474 | 12 |
| rare public (2 ~ 4) | 432 | 23 |
| private TRD | 2617 | 6264 |

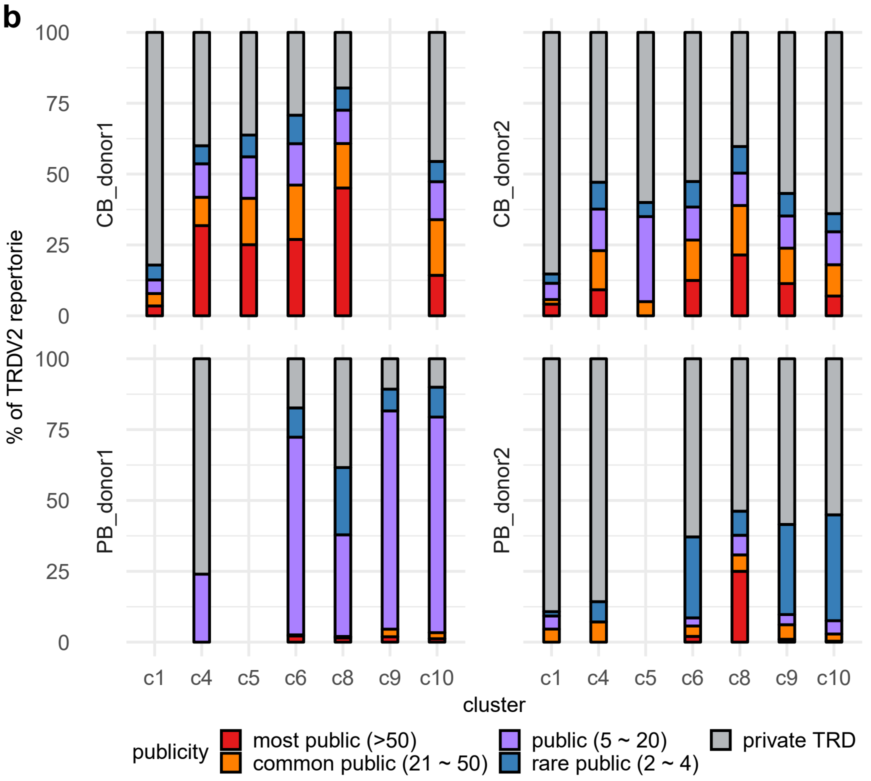


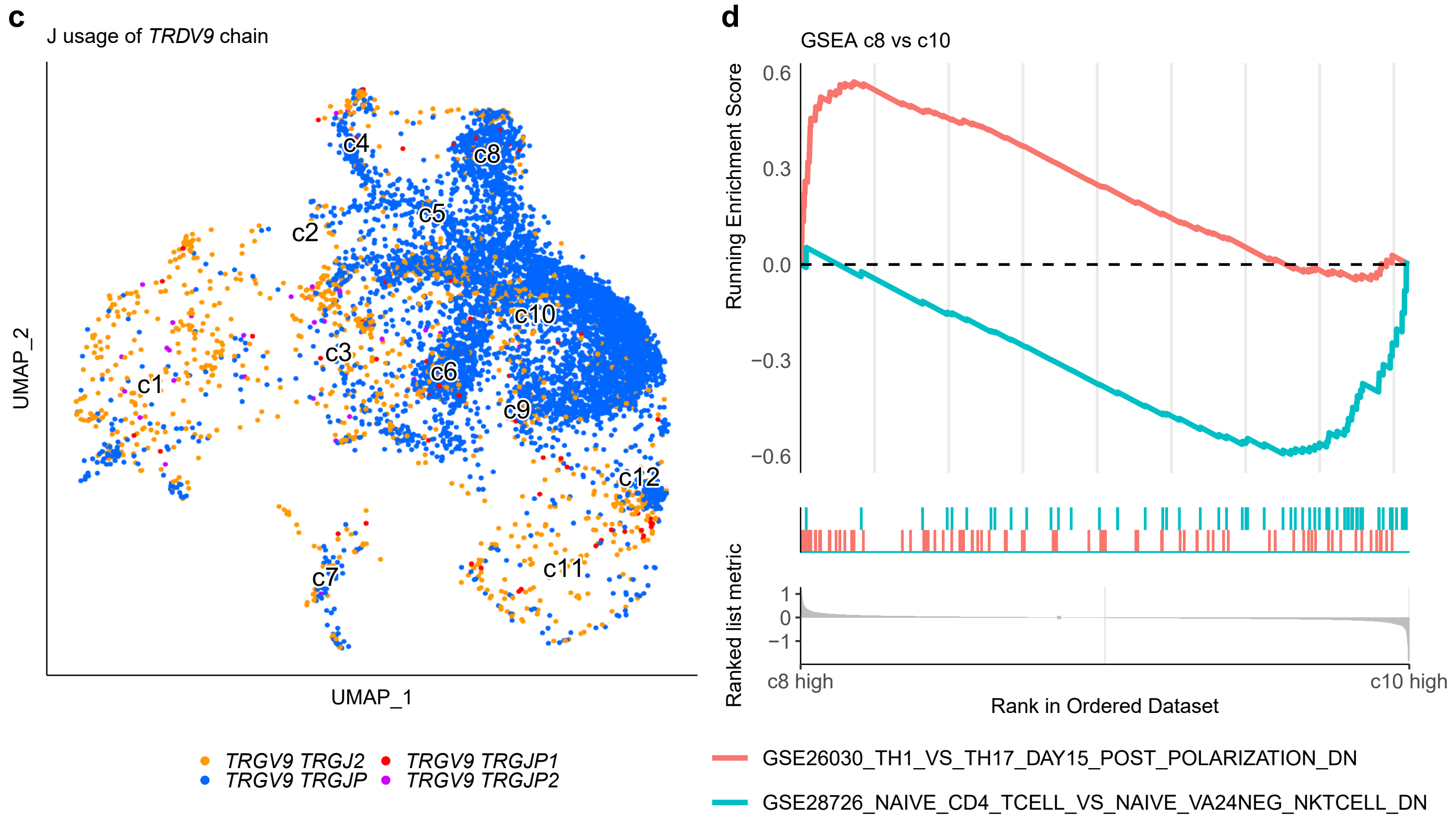


**Figure S5: Vδ2⁺ γδ T cells are heterogeneous innate T cells.**
(**a**) Public *TRD* clones were defined by comparing 80 bulk *TRD* repertoires from independent study populations (ranging from newborns to adults) and four scTCR-seq repertoires in this study. The publicity of different TCRδ chains is summarized in the table. Most public: clones detected in more than 50 donors. Public: clones detected in 21 to 50 donors. Rare public: clones detected in 2 to 4 donors. Private: clones detected in only 1 donor. Only *TRDV2* clones were selected for comparison of scTCR-seq data in **Figure 5a**. (**b**) The publicity of Vδ2⁺ *TRD* clones in the scTCR-seq data is delineated by donor and cluster. (**c**) UMAP of Vγ9⁺ γδ T cells were colored by the J usage of TCRγ chain. (**d**) Gene Set Enrichment Analysis (GSEA) was performed on expression profiles of c8 versus c10 referring to the Immunological gene sets of MSigDb (www.gsea-msigdb.org).

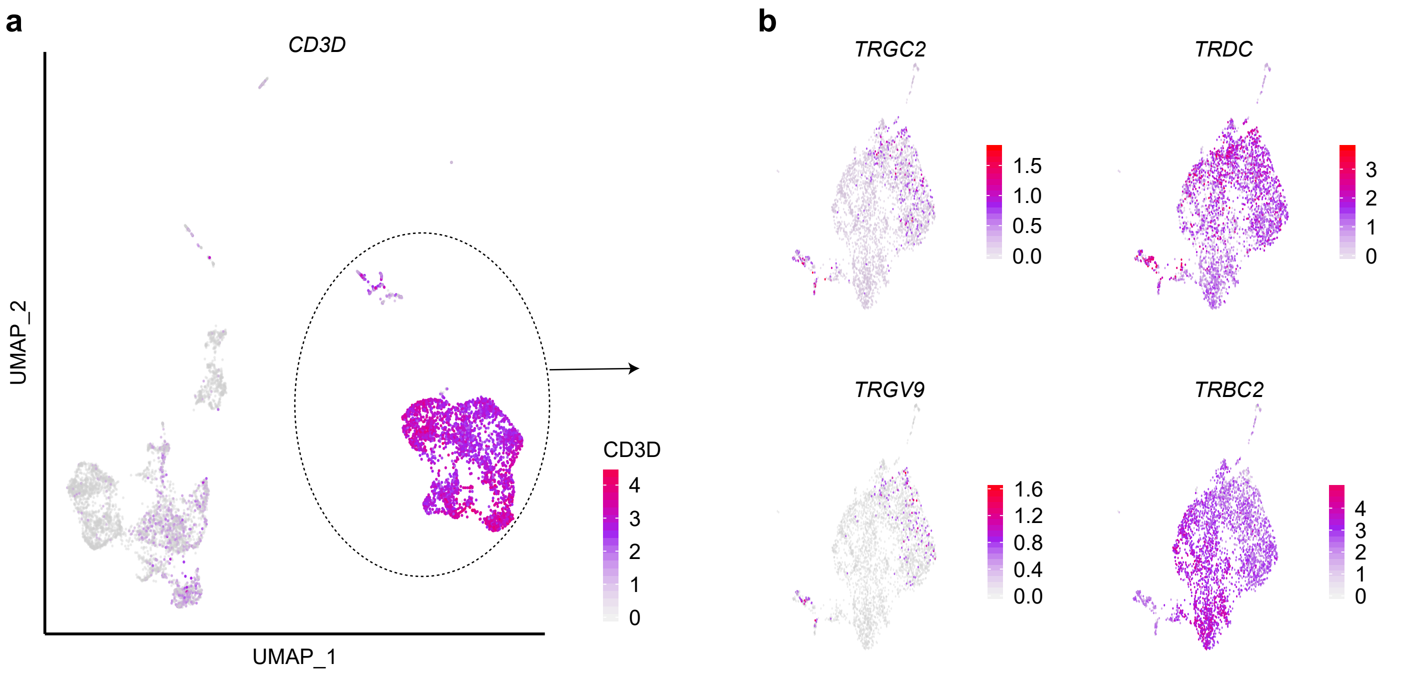


**Figure S6: TCR expression of fetal thymocytes**.
(**a**) Single-cell transcriptome data of human fetal thymocytes from 8 - 10 weeks old embryos (GEO: GSE133341) were subjected to unsupervised clustering and visualized as UMAP. Gene expression of *CD3D* on total fetal thymocytes is shown, while the circle represents selected fetal-thymus T cells for re-analysis in the main **Figure 6**. (**b**) Expression levels of *TRGC2, TRDC, TRGV9*, and *TRBC2* genes on T cells.

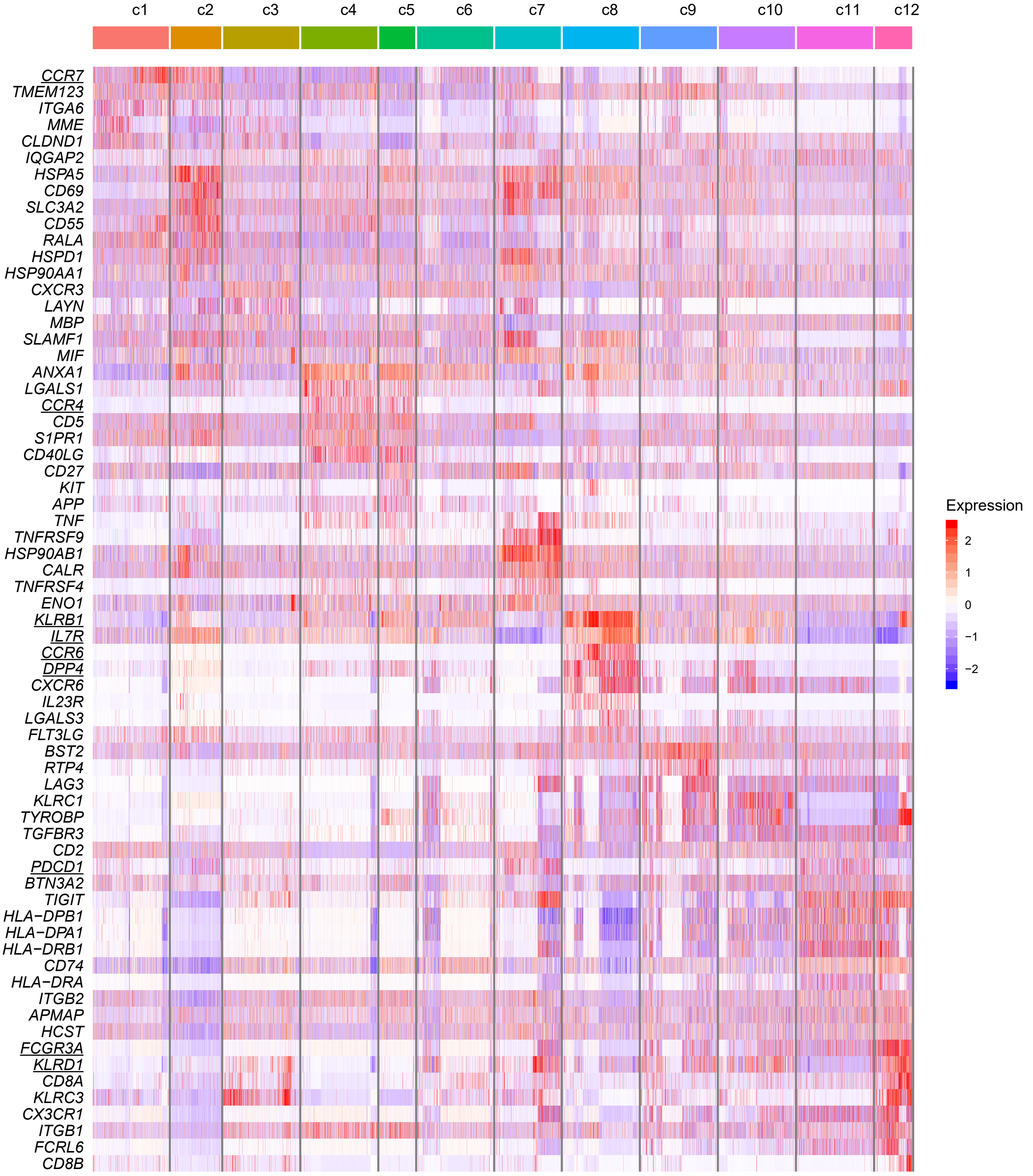


**Figure S7: Differentially expressed surface protein genes.**

Heat map shows Top-10 upregulated surface protein differentially expressed genes (DEGs). The gene list of surface proteins was obtained from GO term cell surface (GO:0009986).

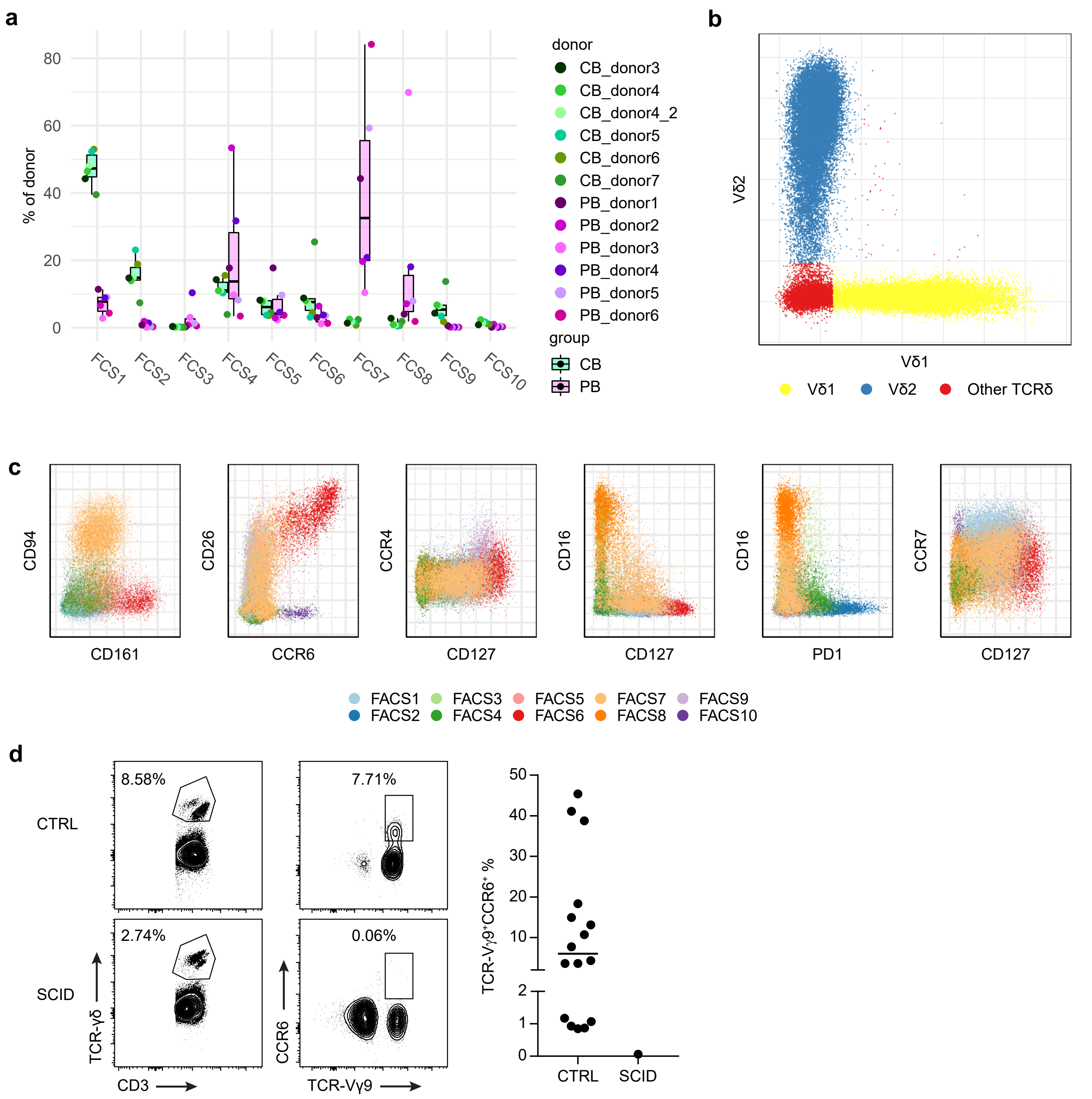


**Figure S8: Multi-color FACS reproduced the heterogeneity revealed by scRNA-seq.**
Samples from five neonates and six adults were tested by multi-color FACS. PB_donor1 and 2 were tested in the scRNA-seq experiment above. Cells from CB_donor4 were tested twice in two independent experiments as an internal control. The TCRγδ$^{+}$ population was pre-gated for bioinformatics analysis. Raw fluorescent values underwent bi-exponential transformation. (**a**) Box-whisker plot highlights the cluster composition in each donor. (**b**) Gating strategy to select Vδ1$^{+}$ and Vδ2$^{+}$ T cells. (**c**) FACS plots of lineage marker expressions on γδ T cells. Cells are colored and grouped by clusters defined in Fig. 7a. (**d**) FACS analysis of a formerly bone marrow-transplanted SCID patient and age- and ex-matched control donor (left panel) and quantification (right panel) of Vγ9$^{+}$ CCR6$^{+}$ γδ T cell frequencies in healthy adults (CTRL, n=16). Each dot represents one donor and the median is shown as a black line.
