## Supplementary material for "A fetal wave of human type-3 γδ T cells with restricted TCR diversity persists into adulthood": methods

### Patient samples and isolation of mononuclear blood cells

Inclusion of healthy donors (n=21), patients (n=1) and cord blood donors (n=7) in this study was performed in accordance with the declaration of Helsinki and approved by the institutional ethics review board at Hannover Medical School under study number 1303-2012 (cord blood donors), 7600-2017 (patient) and 7901_BO_K_2018 (adult healthy donors). All patients gave written informed consent before sample collection. Peripheral blood mononuclear cells (PBMCs) and cord blood mononuclear cells (CBMCs) were isolated from fresh EDTA blood samples using a density gradient media (Pancoll, PAN-Biotech). After isolation, mononuclear cells were washed twice in PBS 10% FBS (Sigma-Aldrich) and cryopreserved at -80°C in 50% FBS (Sigma-Aldrich), 40% RPMI 1640 (Gibco) and 10% DMSO (Roth).

### Single-cell RNA-seq and single-cell TCR-seq libraries

Thawed PBMCs (n = 2) and CBMCs (n=2) were cultured overnight in X-Vivo 15 medium (Lonza), following treatment with 5% Fc-receptor block in PBS (3% FCS; 4 mM EDTA) and antibody staining with hCD3 FITC (1:100; clone BW264/56; Miltenyi) and hTCR γδ PE (1:50; clone 11F2; Miltenyi). DAPI (0.1 µg/ml) was used to discriminate dead cells. γδ T cells were FACS-sorted at the Aria Fusion cytometer (BD). Next, scRNA-seq libraries were prepared from 2-3*$10^4$ FACS-sorted γδ T cells using the Chromium Single Cell 5’ Library Gel Bead and Construction kit (10x Genomics) according to the user guidelines. For generation of single-cell γδTCR-seq libraries, γδ TCR chains were amplified according to the Target Enrichment protocol from cDNA (Chromium Single Cell V(D)J Enrichment Kit, User guide CG000086 Rev J, Step 4) using four reverse custom primers that target the constant region of the *TRG* and *TRD* chain together with the respective forward primers of the kit. In particular, for the first T cell enrichment step the custom primers RV_TRG1: ATCCCAGAATCGTGTTGCTC and RV_TRD1: CCCACTGGGAGAGATGACAA and for the second target enrichment the custom primers RV_TRG2: GGGGAAACATCTGCATCAAG and RV_TRD2: GACAAAAACGGATGGTTTGG were used. Agilent Bioanalyzer High Sensitivity chips were applied for quality control of scRNA-seq and scTCR-seq libraries.

**Next-generation sequencing**

The scRNA-seq libraries were sequenced on the Illumina NextSeq 500/550 platform and the scTCR-seq libraries were sequenced on the Illumina MiSeq or the Illumina NextSeq500/550 platform using read lengths recommended by 10x Genomics.

**Data processing of single-cell RNA sequencing libraries**

The scRNA-seq reads were aligned to the reference human genome GRCh38 (UCUS), following generation of cell-gene matrices via Cell Ranger v3.1 (10x Genomics). Data from ambient RNA was filtered based on nUMI-barcode saturation curve.

**Cell clustering and differentially expressed gene profile**

The R package Seurat v3.1 was used under R v3.6.3 for scRNA-seq data analysis[51,52]. Briefly, cell-gene matrices were imported to R as Seurat projects. Cells contain 1,500 to 6,500 unique molecular identifiers (UMIs) and ≤11% mitochondrial genes were kept for downstream analysis. Cell cycling scores and cell phases were assigned by the function ‘CellCycleScoring’ according to a gene list of cell cycle markers[53]. Raw data matrices were then normalized and scaled by regularized negative binomial regression by ‘SCTransform’ function[54]; contents of mitochondrial genes and ribosomal genesand cycling scores were regressed out as unwanted heterogeneity during normalization. Datasets of each respective sample (n=4) were integrated by Seurat data integration pipeline [51], 2,800 variable genes represent the correspondences between cells in two groups were used as ‘anchors’ to harmonize and integrate the datasets. Integrated data was subjected to principal component analysis (PCA) based on non-mitochondrial/TCR variable genes and 100 principal component (PC) were calculated. The significance of each PC was evaluated by the Jackstraw score. The significant PC were then selected for K-nearest neighbor (KNN) based unsupervised clustering with resolution = 0.8, and Uniform Manifold Approximation and Projection (UMAP) visualization. Finally, obtained clusters with no *CD3* expression (*CD3D*, *CD3E*) were trimmed as contamination.

Clusters defined by unsupervised-clustering were further adjusted based on the number of differentially expressed genes (DEGs) between two neighboring clusters: neighboring DEGs were identified by comparing expression in cells in one cluster to cells in the other cluster by “FindMarkers” function in Seurat. Bimod method was adopted. Neighboring DEGs were defined as average logFC ≥ 0.5 and adjust p-value (based on bonferroni correction) ≤ 0.01. Two neighboring clusters having less than 30 non-mitochondrial/ribosomal DEGs were merged as one cluster.

DEGs among the whole dataset were identified by comparing expression in cells in one cluster to all other cells by "FindAllMarkers" function in Seurat. Bimod method was adopted. DEGs were defined as average log2-fold change (logFC) ≥ 0.25, expressed in minimal percentage cells in at least 1 test group (min.pct) ≥ 15%, and adjust p-value (based on bonferroni correction) ≤ 0.01.

Gene Set Enrichment Analysis (GSEA) was performed and visualized by the R package 'clusterProfiler'[55]. Gene list is arranged by logFC (from high to low) between two clusters and genes expressed in less than 10% of cells within each cluster were excluded. Gene symbols were translated to entrezID. Immunologic gene sets from the Molecular Signatures Database (MSigDB; Broad Institute) were used as reference datasets for GSEA. GSEA was run under 10,000 permutations and the p-value was adjusted by Benjamini-Hochberg method.

**Identification of gene co-expression modules**

DEGs in our neonatal and adult integrated dataset were used for the identification of gene co-expression modules (GMs). After excluding all mitochondrial and ribosomal DEGs, the average expression values of DEGs in each cluster were calculated, log-transformed and scaled. The scaled matrix was embedded by UMAP for visualization. GMs were identified by hierarchical clustering method under 'complete' model (height cutoff = 5.7). Clusters with similar expression patterns were merged as one GM. Gene ontology (biological process) enrichment analysis was performed on GMs by the R package 'clusterProfiler'. Aggregated GM expression scores were calculated on single-cell base by 'AddModuleScore' function from Seurat.

**Single-cell TCR-seq analysis**

Cell Ranger VDJ pipeline (10x Genomics) under default parameters was performed to generate scTCR annotations on the assembled contigs and the clonotype consensus sequences from fastq files. Human genome GRCh38 was used as reference for alignment. The scTCR-seq annotation tables from multiple sequencing runs from the same sample were merged. Non-productive TCR sequences as well as duplicates (i.e. one barcode has two *TRD* or *TRG* reads) were excluded. The merged annotation tables were exported as processed data. Further, by matching the barcodes, scTCR-seq data was incorporated into the metadata of above created scRNA-seq Seurat project. TRG and TRD clones were defined based on amino-acid sequences. Gini indices of paired TCR clones were calculated by R package reldist.

Public *TRD* clones were defined based on comparison of single-cell *TRD* repertories from four donors in this study and bulk *TRD* repertories from 80 donors[9,16,56] (and Ravens et al. accepted, SRA accession code PRJNA592548). Among them were 21 adults and 63 children aged between 0 and 3.5 years. CDR3 regions were used for comparison. One *TRD* clone was defined as public clone if its CDR3 sequence was recovered from at least two donors. According to the number of donors that share this clone, we classified the publicity of public clones as rare public (2 to 4 donors), public (5 to 20 donors), common public (21 to 50 donors), and most public (more than 50 donors).

**Single-cell transcriptome analysis of fetal thymocytes**

Embryonic week 8-10 thymocytes expression matrices were acquired from GEO: GSE133341 and analyzed as described above for scRNA-seq data. In brief, cells containing 2,500 to 20,000 UMIs and ≤0.5% mitochondrial genes were considered for analysis. After dimensional reduction and clustering on all thymocytes, the clusters with high *CD3D* expression were selected as the T cell subset. Identified T cells were subjected to another round of dimensional reduction analysis and clustering. Cluster identity was assigned by the expression of TCR genes. GM scores were calculated based on GMs identified in the two cord blood and two adult peripheral blood datasets of this study.

**Multi-color flow cytometry**

Flow cytometry was performed using a LSRII cytometer (BD) and cell suspensions of thawed PBMCs (n = 6) or CBMCs (n = 5) ($2*10^6$ cells) were treated with 5% FcR block in buffer (PBS with 3% FCS, 4 mM EDTA) before staining. The following antibodies were used: hTCR γδ PE (1:50; clone 11F2; Miltenyi), hTCR Vδ1 VioGreen (1:50; clone REA173; Miltenyi), hTCR Vδ2 PerCP-Vio700 (1:50; clone REA771; Miltenyi), hCD26 APC (1:50; clone BA5b; BioLegend), hCD94 PE-Cy7 (1:50; clone DX22; BioLegend), hCCR4 BV650 (1:50; clone 1G1; BD), hCCR6 BV785 (1:50; clone G034E3; BioLegend), hCCR7 BV711 (1:25; clone G043H7; BioLegend), hPD1 BV605 (1:20; clone EH12.2H7; BioLegend), hCD16 FITC (1:200; clone 3G8; BioLegend), hCD161 APC-Cy7 (1:25; clone HP-3G10; BioLegend) and hCD127

BV421 (1:25; clone A019D5; BioLegend). DAPI (0.1 μg/ml) staining was used in a separate control to determine living cells.

Whole blood of healthy controls and a SCID patient was analyzed as previously described (https://doi.org/10.1101/2020.05.11.20096263). Briefly, 500μL of whole blood was processed in erythrocyte lysis buffer. Samples were stained for 20 mins in PBS at RT, washed twice and acquired on a Cytek Aurora spectral flow cytometer. Cytometry data were analyzed using FlowJo v10 software (Tree Star). The following antibodies were used: Zombie NIR (Biolegend), hCCR6 BV711 (Biolegend), hTCR Vγ9 BV605 (BD), hTCR γδ APC (Miltenyi) and hCD3 AF532 (Invitrogen).

**Multi-color flow cytometry data analysis**

Flow cytometry (FACS) data was analyzed based on the CyTOF workflow[57]. Briefly, TCRγδ$^+$ subsets from PBMC and CBMC were gated and isolated by FlowJo software (BD) and exported to R v3.6.3. Biexponential transformation was applied to raw fluorescent intensities allowing to remove background fluorescence and spreading error[58]. Scaled expression values of nine lineage markers (CCR6, CD26, CD94, CD161, CD16, CD26, CCR4, CCR7, and PD1) were subjected for un-supervised clustering and UMAP embedding. Un-supervised clustering was conducted with R package “FlowSOM” [59] and “ConsensusClusterPlus” [60]. Firstly, a self-organizing map (SOM) was built to enwrap all cells into 100 SOM nodes. “ConsensusClusterPlus” was then called on SOM nodes using hierarchical clustering method. The number of clusters was firstly determined by Empirical cumulative distribution function (CDF) plots and afterwards clusters with similar expression profiles were merged together.

**Data availability**

Raw data and codes for analysis can be acquired from corresponding author upon to request.
